# MORPHE: Bridging Image Generation and Spatial Omics for Tissue Synthesis

**DOI:** 10.64898/2026.03.03.709377

**Authors:** Yuan Feng, Zachary Robers, Leyla Rasheed, Yang Miao, Shuo Wen, Kevin Lee, James Sohigian, Maria Brbić, John W. Hickey

**Author notes:** Contributing authors.

## Abstract

Spatially resolved omics technologies reveal tissue organization at single-cell resolution but remain limited by the cost of the assays, incomplete spatial coverage, 2D-only imaging, and experimental artifacts. These factors motivate the need for *in silico* methods that can reconstruct or extend tissue context beyond what current spatial measurements provide. We present MORPHE (MOdeling of stRuctured sPatial High-dimensional Embeddings), an AI framework that learns to synthesize biologically faithful tissue architecture directly from spatial-omics data. MORPHE introduces a graph-informed probabilistic embedding that maps discrete cell identities and their spatial relationships into a continuous RGB-like latent space compatible with diffusion modeling. This representational bridge enables spatial cellular maps to leverage large pre-trained image-generative models while preserving biological interpretability upon decoding. By modeling cells as the fundamental units of generation and learning how their identities and spatial relationships collectively give rise to large-scale tissue structure, MORPHE enables generation and reconstruction of tissue architecture at single-cell resolution. We applied the method across large-scale single-cell proteomic datasets from the intestine and single-cell transcriptomic datasets from the brain, showing computational scalability acrosss millions of cells. We used MORPHE on these datasets to outpaint beyond experimentally restricted fields of view, inpaint missing or experimentally damaged tissue regions, and perform cross-tissue imputation, connecting separated tissue regions into a single contiguous sample in both 2D and 3D. MORPHE represents a new class of tissue generation algorithms that will help solve current limitations and challenges with single-cell spatial-omics datasets.

## 1 Introduction

Spatially resolved omics technologies have transformed our ability to interrogate tissues at single-cell resolution. By jointly capturing molecular and spatial information, these approaches have enabled unprecedented insights into how cellular composition and organization govern tissue function in health and disease [1–8]. These efforts have already revealed new organizational principles, ranging from tissue-specific niches to pathological remodeling, and they have been one of the key technologies driving large-scale mapping consortia efforts [9–17].

Despite this progress, spatial omics remains constrained by technical and experimental challenges. Laboratory assays are constrained by cost and sample availability (e.g., patient biopsy size), restricting measurements to limited tissue regions and very limited number of 3 dimensional (3D) datasets. Thus, many spatial datasets only provide fragments rather than continuous or full tissue landscapes, which can hamper cross-sample comparisons. Moreover, tissue processing and imaging can introduce artifacts that compromise analyses (e.g., rips, autofluorescent fibers, and bubbles). Datasets containing truncations, gaps, or missing areas pose problems for many downstream analyses that require intact, contiguous tissue structure, such as cell-cell interaction analyses, neighborhood identification, and enrichment analyses [3, 4, 9, 11, 18]. Therefore, there is a critical need for generative approaches that can reconstruct or extend tissue architecture to recover missing tissue areas, extend areas not imaged, and interpolate between discontinuous tissues in multiple dimensions.

Recent work in spatial omics has introduced a range of computational approaches for inferring tissue organization from molecular measurements. Reference-based mapping methods such as Tangram [19], CytoSPACE [20], and CeLEry [21] project dissociated cells onto predefined spatial atlases, but rely on explicit reference tissues and cannot synthesize new spatial regions. De novo reconstruction approaches, including novoSpaRc [22], infer spatial layouts directly from transcriptional similarity without a predefined atlas, yet still require gene expression measurements from the regions being reconstructed. LUNA [23] learns atlas-derived spatial priors to reassemble dissociated cells but cannot generate new cell states or molecular profiles in unmeasured regions. MIMYR [24] introduces a multi-stage generative framework that reconstructs missing spatial transcriptomic slices by sequentially modeling cell locations, cell identities, and gene expression profiles; yet this only generates over spatial transcriptomics. SpatialZ [25] aims to bridge planar spatial transcriptomics slices into threedimensional atlases, but primarily performs interpolation across adjacent sections through alignment and lookup-based assignment rather than generative imputation of missing tissue.

To create a broadly applicable generative framework operating solely on spatial-omics tissue maps, we introduce MORPHE (MOdeling of stRuctured sPatial High-dimensional Embeddings), a generative framework that models tissue organization as a diffusion process over structured cellular representations. MORPHE represents single-cell–resolution tissue maps in a continuous, image-like representation by encoding graph-informed probabilistic cellular information into a 3D continuous space. This allows the framework to leverage priors from large-scale pretrained image generative models that exist in a 3D RGB space. Moreover, a cascaded diffusion architecture is used to further refine generated regions and achieve pixel-level precision for single-cell resolution. This formulation enables MORPHE to coherently generate, extend, and reconstruct tissue regions at single-cell resolution, establishing it as a general computational framework for spatial generation that can be applied across diverse spatial-omics datasets (e.g., transcriptomic and proteomic data).

We evaluated MORPHE on multiple large-scale spatial-omics datasets generated using different technologies and marker panels, including a 2.6 million–cell Co-detection by indexing (CODEX) dataset of the human intestine [11], a 2.1 million–cell Multiplexed Error-Robust Fluorescence In Situ Hybridization (MERFISH) mouse dataset [9], and a 6.1 million–cell whole MERFISH mouse brain atlas composed of serial tissue sections [10]. Across these datasets, MORPHE consistently synthesized tissue regions with high structural, compositional, and cellular neighborhood fidelity, demonstrating robust performance across distinct spatial-omics modalities, marker panels, and tissue-specific spatial organizations.

Within this unified framework, we showed that MORPHE supports four complementary applications that address common experimental limitations of spatial omics technologies. First, MORPHE can inpaint missing or experimentally damaged tissue regions, enabling virtual restoration of disrupted neighborhoods. Second, MORPHE can outpaint beyond restricted fields of view, allowing extension of partial tissue samples. Third, MORPHE can perform 2D cross-tissue imputation, which connects spatially separated tissue regions within a plane into coherent structures. Fourth, MORPHE can perform 3D cross-tissue imputation, in which spatial information from two non-adjacent tissue sections conditions the generation of volumetric tissue structure connecting them across depth. Together, these capabilities transform fragmented spatial measurements into continuous tissue landscapes, facilitating downstream spatial analyses, improving cross-sample comparability, and enabling *in silico* exploration of tissue organization beyond existing datasets.

## 2 Results

### 2.1 MORPHE Bridges Spatial-omics and Diffusion-based Image Generation

Generative models, especially image-based generative models, have demonstrated remarkable capabilities in image synthesis. In recent years, increasingly mature image generation approaches have been able to produce high-quality, sharp images with clear structural and semantic coherence. To test if we could leverage pretrained image-based models to provide a generative framework to spatial-omics data, we used stable diffusion 2 (SD2) [26] on a cell type map of a spatial proteomic dataset of the healthy human intestine[11] generated a rough approximation of color and some structure (**Extended Data Fig. 1a**). However, the generation was both inaccurate and uninterpretable because there are not discrete boundaries between cells and there is blending the colors of cell types that does not resemble the original structure (**Fig. 1a**). Thus, to make spatial-omics data compatible with off-the-shelf image generation architectures [26–29], we needed to solve two challenges: physical separation of cell types in spatial images and transforming the color dimensional space into one that encodes biologic meaning.

**Fig. 1.**
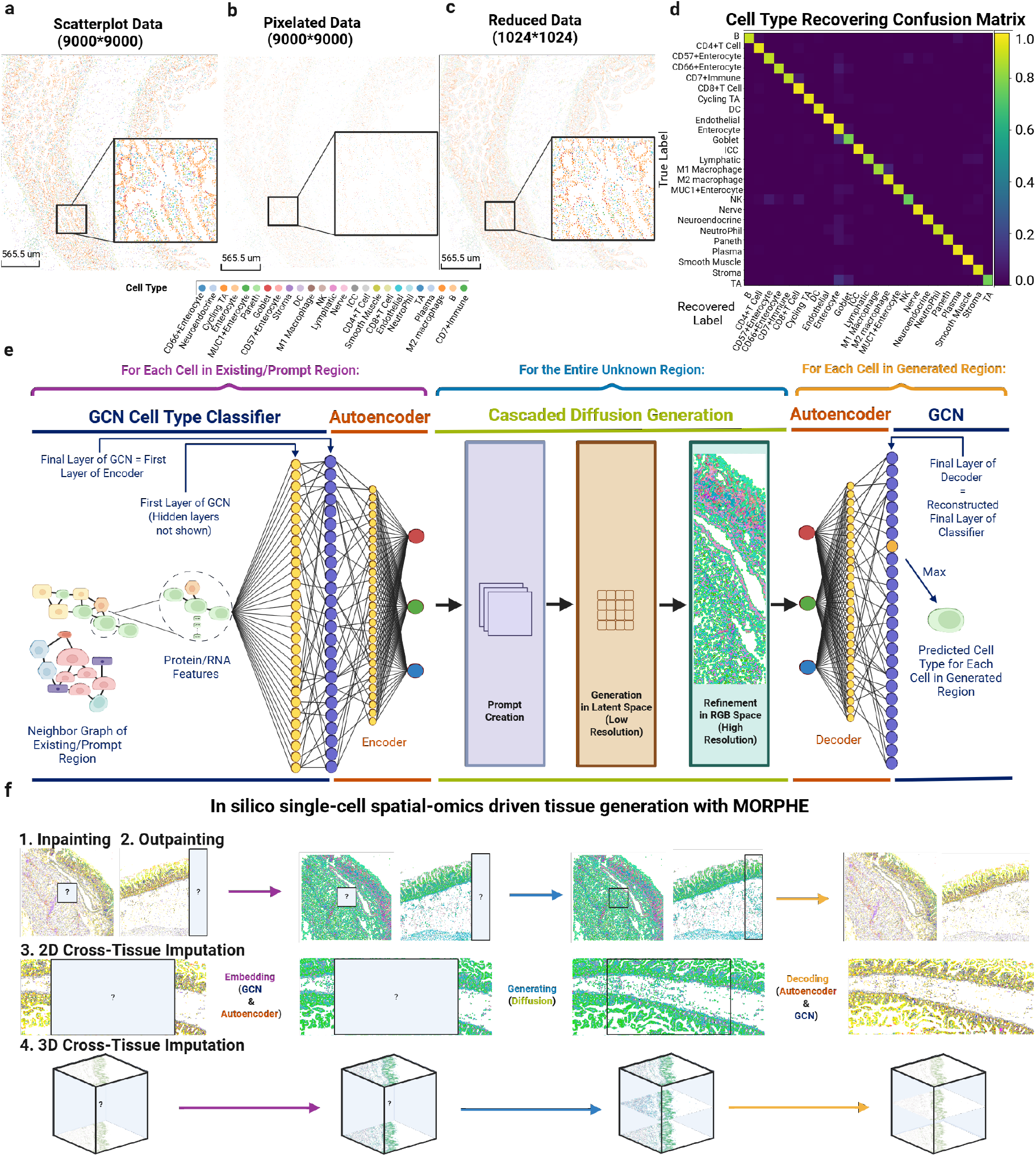
MORPHE is a generative framework that models tissue organization as a diffusion process over structured cellular representations. **a)** An image of the original dataset at 9000×9000 resolution, with each cell represented by a large cluster of pixels. **b)** An image of the 9000×9000 dataset with each cell represented by a single pixel. **c)** An image of the reduced 1024×1024 dataset with each cell represented by a single pixel. **d)** Confusion matrix for classification accuracy across cell types in the entire CODEX intestine dataset. **e)** At the single-cell level, MORPHE uses a GCN that operates on cell-level k-nearest neighbor graphs enriched with molecular features (i.e. Protein or RNA). The final GCN layer yields an n-dimensional vector of soft cell-type probabilities for n cell types; these logits serve as the input layer of an autoencoder. The encoder compresses the n-dimensional (n = number of cell types) vectors into a 3-dimensional RGB-like representation that forms the input to the cascaded diffusion process. Generation proceeds by constructing a prompt from the RGB-like embeddings, performing coarse denoising in a latent space, and refining the output in a high-resolution RGB space. The generated RGB embeddings are then decoded back to n-dimensional logits, and the cell type is assigned using the maximum-probability entry. **f)** MORPHE performs tissue generation at the single cell level using single-cell spatial-omic prompts. It supports four main use cases: missing tissue inpainting, truncated tissue outpainting, 2D cross-tissue imputation and 3D cross-tissue imputation. For each use case, MORPHE follows three-stage generative framework: (i) embedding observed cells into a learned 3D RGB-like latent representation, (ii) performing conditional diffusion within this space to generate embeddings for the unknown region, and (iii) decoding the generated embeddings back into cell-type representations. Colors in the intermediate panels denote continuous RGB-like latent embeddings, whereas colors in the first and final panels represent discrete cell types.

First, to discretize cell types in space from spatial-omics datasets we formulated images as cells as colored pixels based on their centroid x, y location (**Fig. 1b**). However, we observed a high number of empty pixels between cells, due to high resolution imaging. To enable training with less compute and focus the model to learn important multicellular interactions and structures, we designed a method to merge nearby pixels into a downscaled grid (**Extended Data Fig. 1b**). Using this method, we observed that we could obtain a denser representation, where we could get 81-fold compression to a 1024×1024 map that retained 99.58% of cells, after which there was a steep loss in cells (**Extended Data Fig. 1c**) [11]. Importantly, while eliminating unnecessary spatial sparsity, the compressed representation preserves global tissue architecture and maintains clearly delineated, pixel-resolved cell boundaries (**Fig. 1c**). Finally, because we still observed significant empty pixels, we further downsampled to 512×512 using nearest-neighbor interpolation which preserved all cells and structural relationships.

Second, we encoded categorical cell-type identities into a continuous RGB-like space. Initially, we attempted assigning fixed RGB values to each of the 25 cell types. However, this introduced sharp color-space discontinuities, resulting in large regions converging to a small number of dominant colors and a loss of spatial diversity and biological accuracy which caused by pattern collapse within the encoder (**Extended Data Fig. 2 a-f)**.

To make this encoding informed by each cell’s local neighborhood context and molecular profile, we extracted the neighborhood-aware soft probability vectors with a Graph Convolutional Network (GCN) (See Section 4.3) [30] (**Extended Data Fig. 1d**). We then used an Autoencoder (See Section 4.4) (**Extended Data Fig. 1e**) to compress these probabilistic cell-type embeddings into a continuous, three-channel latent representation to match the RGB dimensionality of pixels in images (**Extended Data Fig. 1f**). This transformation enables us to leverage large-scale pretrained image generative models [26] that allow MORPHE to leverage and finetune its learned priors, while retaining biologically meaningful cell-type relationships.

To evaluate whether the RGB-like embedding faithfully preserves cellular identity prior to any generative modeling, we projected cells into RGB-like latent space using neighborhood-informed features and then reconstructed their original cell-type labels from the RGB-like representation. We quantified recovery accuracy across all regions of a healthy human intestine spatial proteomic dataset comprising 25 cell types, 66 tissue regions, and 2.6 million cells [11]. The embedding achieves an average reconstruction test-set accuracy of 92.5%, with 23 of 25 cell types exceeding 90% average recovery accuracy (**Extended Data Fig. 1g**).

For the two lower-performing cell types, misclassifications remain biologically coherent. Goblet cells are primarily confused with enterocytes, both epithelial subtypes, as shown in the confusion matrix (**Fig. 1d**). Similarly, transit-amplifying (TA) cells—partially differentiated epithelial cells—are most frequently misclassified as enterocytes or goblet cells. These structured and biologically plausible confusions indicate that the embedding preserves meaningful cellular similarity relationships as these share many of the same protein markers. By encoding probabilistic neighborhood-informed identities into a continuous RGB space, MORPHE maintains biological interpretability even when minor semantic shifts occur, constraining errors within related cell lineages rather than across unrelated cell classes.

Additional genomic or proteomic features can further improve the reconstruction accuracy. In the case of a MERFISH mouse cortex dataset consisting of 58 tissue regions and approximately 2.1 million cells [31], the model achieved 99.05% testset-accuracy. When trained on a coronal MERFISH 3D whole mouse brain datasets comprising 183 sections and approximately 6.1 million cells [10], it demonstrated 96.02% testset-accuracy. These results demonstrate that we bridge the gap between the traditional representation of spatial omics datasets and image representations of tissue regions by establishing a complete pipeline that transforms spatial omics data into continuous, interpretable, image-like representations suitable for generative modeling.

With this accuracy, we then designed a cascaded diffusion architecture to perform spatial generation in three coordinated stages (**Fig. 1e, Extended Data Fig. 1h-k**). First, the visible tissue prompt is transformed to RGB-like embeddings using the GCN and autoencoder and are mapped through a convolutional encoder to extract local morphological features. These are fused with positional embeddings of the masked coordinates to form a high-dimensional prompt that encodes spatial organization and cellular composition of the known region (**Extended Data Fig. 1i**). Second, conditioned on this prompt, the latent diffusion model which we take advantage of the large-scale pretrained image generation model [26] denoises a low-resolution latent representation of the missing region, akin to as shown in **Extended Data Fig. 1j** [29]. Iterative cross-attention and convolutional blocks allow the model to propagate long-range spatial dependencies while generating coarse but consistent global structure. Third, the output from the latent space generation is decoded into an RGB-like embedding map by a second diffusion stage that restores cell-scale structure (**Extended Data Fig. 1k**). This refinement module outperforms existing VAE-based decoders, ensuring semantic consistency at the cellular level while preserving high pixel-level fidelity (**Extended Data Fig. 2g**).

Overall, by unifying graph-based cell representation learning with cascaded diffusion generation, MORPHE establishes a biologically grounded yet image-compatible generative framework for spatialomics data. Rather than directly modeling discrete cell labels, it operates in a continuous probabilistic embedding space that preserves uncertainty and cell-type relationships. This design enables the model to leverage large-scale pretrained image priors to support: Inpainting of missing regions, outpainting beyond limited fields of view, and both 2D and 3D cross-tissue imputation depending on the provided spatial prompts (**Fig. 1f**). The refinement diffusion process separates global structural reasoning from fine-scale morphological refinement, allowing MORPHE to generate tissues that are coherent at both architectural and cellular resolutions.

### 2.2 MORPHE Outpaints Tissue Regions Beyond Observed Boundaries

Our first application of MORPHE is outpainting that extends spatially resolved tissue maps beyond their captured boundaries, enabling the *in silico* reconstruction of broader tissue contexts. Using RGB-embedded representations, MORPHE iteratively generates new tissue patches by concatenating predicted areas while maintaining a fixed input window; the generated latent features are then refined and decoded by the diffusion decoder to produce final single-cell–resolution tissue extensions (**Fig. 2a**). This solves a critical problem because each spatial-omics dataset remains a finite and incomplete snapshot of biological structure coming from limitations in imaging field-of-view, tissue sectioning, and biopsy sampling that truncate the native spatial context, obscuring large-scale organizational patterns and interactions across regions.

**Fig. 2.**
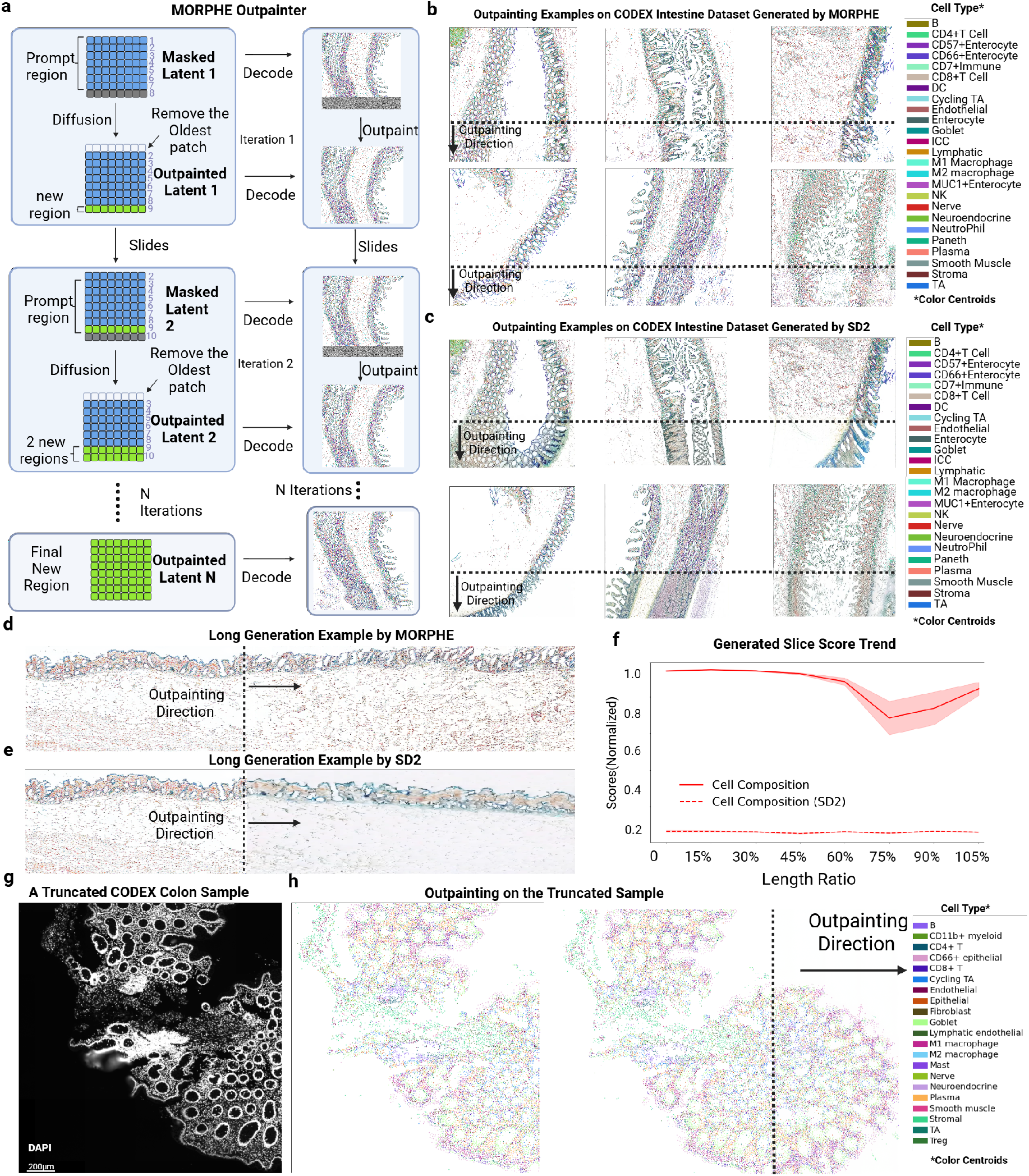
MORPHE extends spatially resolved tissue maps beyond captured regions, generating coherent and biologically consistent structures through diffusion-based outpainting. **a)** MORPHE’s outpainter iteratively generates and concatenates new regions and removes the furthest row from the newly concatenated regions until N new rows are generated to extend the region. **b)** Multiple outpainted examples from CODEX dataset generated by MORPHE. **c)** Multiple outpainted examples from CODEX dataset generated by Stable Diffusion 2 (SD2). **d)** An example of MORPHE outpainting up to 105% times the size of the original region and **e)** SD2 for the same region as **d). f)** Outpainting quality of MORPHE&SD2 as it varies with the length of the outpainted region. The line represents the average across ten trials, while the shaded area denotes the range (upper and lower bounds) among them for density, structure, and composition scores. **g)** Example of a CODEX healthy colon sample with a region truncated during imaging. The rightmost portion of the tissue is entirely missing in the original acquisition. **h)** Shown are the corresponding pixelated embedding after MORPHE preprocessing for **g)**, and the tissue region outpainted by MORPHE to restore the missing spatial context.

We apply this first to the healthy human intestine spatial proteomic dataset. We chose this for four reasons: first, it has natural and previously characterized structures; second, it has diverse structures in the same image (e.g., mucosa, submucosa, and muscularis externa); third, different cross sectional slices were taken; and fourth, we have expertise in the biological organizations. We further verified across multiple complementary metrics that the intestine dataset exhibited substantial inter-region heterogeneity at architectural, perceptual, compositional, and neighborhood scales (**Extended Data Fig. 3**). This provides a rigorous and diverse testbed for evaluating MORPHE’s ability to generalize across distinct tissue architectures.

Applying MORPHE, we see that MORPHE’s performance generalizes across multiple intestinal regions from our test dataset (**Fig. 2b**). Indeed it consistently produces structurally coherent and semantically plausible tissue, outperforming Stable Diffusion 2 (SD2) [26] (**Fig. 2c**), a popular outof-the-box image outpainting method, which we replace our cascaded diffusion module by it and apply the embedding module on it for comparison. Moreover, in a representative transverse sample, MORPHE maintained structural continuity and biologically plausible organization across even further extended regions (**Fig. 2d**), whereas SD2 exhibited earlier fragmentation (**Fig. 2e**).

To assess the stability and generative limits of MORPHE, we evaluated long-range generation using cell composition similarity (Jensen–Shannon divergence of cell-type distributions) [32]. Across ten independent trials extending tissues to 1.05× their original length (**Fig. 2f**), MORPHE outperformed SD2 across nearly all evaluated extension lengths on overall biological consistency. Although SD2’s performance appeared more stable with distance, its overall score remained consistently low across the entire generation range, especially the cell composition similarity, indicating an inability to preserve global cell-type distributions during extrapolation. In contrast, MORPHE’s fidelity gradually declined yet stabilized beyond approximately 75% of the generated length while maintaining substantially higher overall biological coherence.

Observing high quality generation, we used MORPHE to outpaint a healthy colon sample from a different spatial proteomics dataset of the intestine. The rightmost region was partially truncated during imaging, due to issues with tissue placement on the slide (**Fig. 2g**). MORPHE restored the missing spatial context based solely on information from the observed tissue (**Fig. 2h**).

At single-cell resolution, we examined the spatial distribution of specific immune populations, with a particular focus on M1 macrophages in the colon. These cells are known to preferentially localize near the lumen, where they contribute to immune surveillance and pro-inflammatory responses against microbial invasion or tissue damage. Notably, the generated tissue recapitulates this characteristic pattern, with M1 macrophages enriched along the tissue boundary and relatively depleted in central regions, closely matching their distribution in the original tissue (**Extended Data Fig. 4a**). As a comparison, we also visualized the spatial distribution of M2 macrophages, which exhibited a more uniform distribution across the tissue in both the original and generated samples (**Extended Data Fig. 4a**).

Spatially, it is clear that separate intestinal crypt structures and the interdispersed stromal and immune contexts are preserved recovering lost tissue, consistent with what we observed in the unannotated, truncated raw H&E images (**Extended Data Fig. 4b**). While we do not have ground truth for this dataset, we also evaluated its output by performing multicellular neighborhood analysis on the reconstructed region (**Extended Data Fig. 4c**). We observed that the outpainted area preserves neighborhood organization that closely matches the original tissue, such as the characteristic concentric arrangement of goblet-enriched and epithelial-enriched neighborhoods (**Extended Data Fig. 4d**).

At a higher-order biological scale (clustering of neighborhoods) as defined in **Extended Data Fig. 4b**, we quantitatively observe layering of 5 spatial categories (**Extended Data Fig. 4e**). This again recapitulates the correct layering of the intestine in an inside-outside layering which is based on higher-order multicellular structures across the intestine (**Extended Data Fig. 4f**). Without out-painting, truncation of the tissue leads to an overestimation of the bottom layer and a corresponding underestimation of the top layer (**Extended Data Fig. 4g**). Consequently, any analysis relying on architectural layer percentages would be systematically biased by tissue incompleteness. This shows that MORPHE restores multiscale biological structures from individual cell type densities to cellular neighborhoods to higher-scale tissue unit level organization.

Taken together, these results suggest that MORPHE enables biologically coherent and spatially consistent tissue outpainting beyond experimentally captured regions. Through its cascaded diffusion framework and diffusion-based decoder, MORPHE can extend tissue structures while maintaining global composition and microarchitectural fidelity. This approach offers a practical path toward reconstructing broader tissue contexts *in silico*, helping to bridge the gap between limited imaging fields and the continuous organization of real tissues.

### 2.3 Integrating Established and Novel Spatial Metrics for Comprehensive Evaluation of MORPHE

With biological accuracy and initial evaluations, we moved to more comprehensively compare MORPHE’s performance by performing mulitple replicates, comparing to a number of architectures, comparing to multiple ground truths, and developing new metrics for spatially generated data. We first generated tissue extensions corresponding to 25% of the original tissue length five times per sample. We benchmarked MORPHE against multiple state-of-the-art image-generation algorithms: Stable Diffusion 2 (SD2) [26], FLUX.1 Fill Dev (FF) [33], a commercial image-editing application (Pixelcut) [34]. As shown in **Extended Data Fig. 2c**, applying standard image generation methods without our spatial embedding framework leads to poor performance and does not yield a meaningful comparison. Therefore, to isolate and fairly evaluate the generative capacity of different models, we replaced only MORPHE’s cascaded diffusion module with each baseline method while keeping all other components of the pipeline fixed, including the dimensionality reduction, GCN, and autoencoder. By holding the shared embedding and spatial encoding stages constant, this experimental design ensures that any observed differences in performance arise specifically from the generative model itself rather than from upstream representation learning. All generated outputs are compared against a held-out tissue region from the same dataset.

While comparing to held-out tissue regions is useful, we reasoned that because of natural heterogeneity in tissue composition and organization across a tissue, these comparisons would not account for this natural heterogeneity. For example, if you compare the left half of a tissue to the right half, while there will be substantial similarities, they are not exact copies of each other and vary in both cellular composition and structural location. Consequently, we also split each tissue region into generated part, held-out true part and reference part (**Extended Data Fig. 5a**). The generated part is directly compared against to its adjacent part in the prompt region named “held-out true”, while the held-out true region is compared again to its adjacent part named “reference” to provide a standard, thereby contextualizing quantitative generation scores with respect to the inherent variability present in the original tissue.

First, to evaluate the accuracy of cell-type composition, we computed the KL divergence between the cell type distributions of real and generated regions. MORPHE achieves a mean score 67% higher than competing methods (**Fig. 3a**). In the same duodenum sample, MORPHE’s generation reproduces the original cell type proportions, while SD2 produces excess M2 Macrophages and Paneth cells (**Fig. 3b**), underscoring the need for MORPHE’s cascaded diffusion architecture, which replaces the conventional VAE decoder with a diffusion-based decoder [26]. Unlike VAE decoders that reconstruct pixel-level outputs in a single step—often introducing color drift and blurring—MORPHE’s diffusion decoder progressively refines latent features through iterative denoising, recovering high-frequency spatial details and sharp cellular boundaries. This design enables pixel-level (single-cell–level) precision and preserves local compositional integrity, allowing MORPHE to more faithfully reproduce the true cellular makeup of tissues. Consequently, the diffusion-based decoding step is key to MORPHE’s superior compositional fidelity and its substantial performance boost over competing methods.

**Fig. 3.**
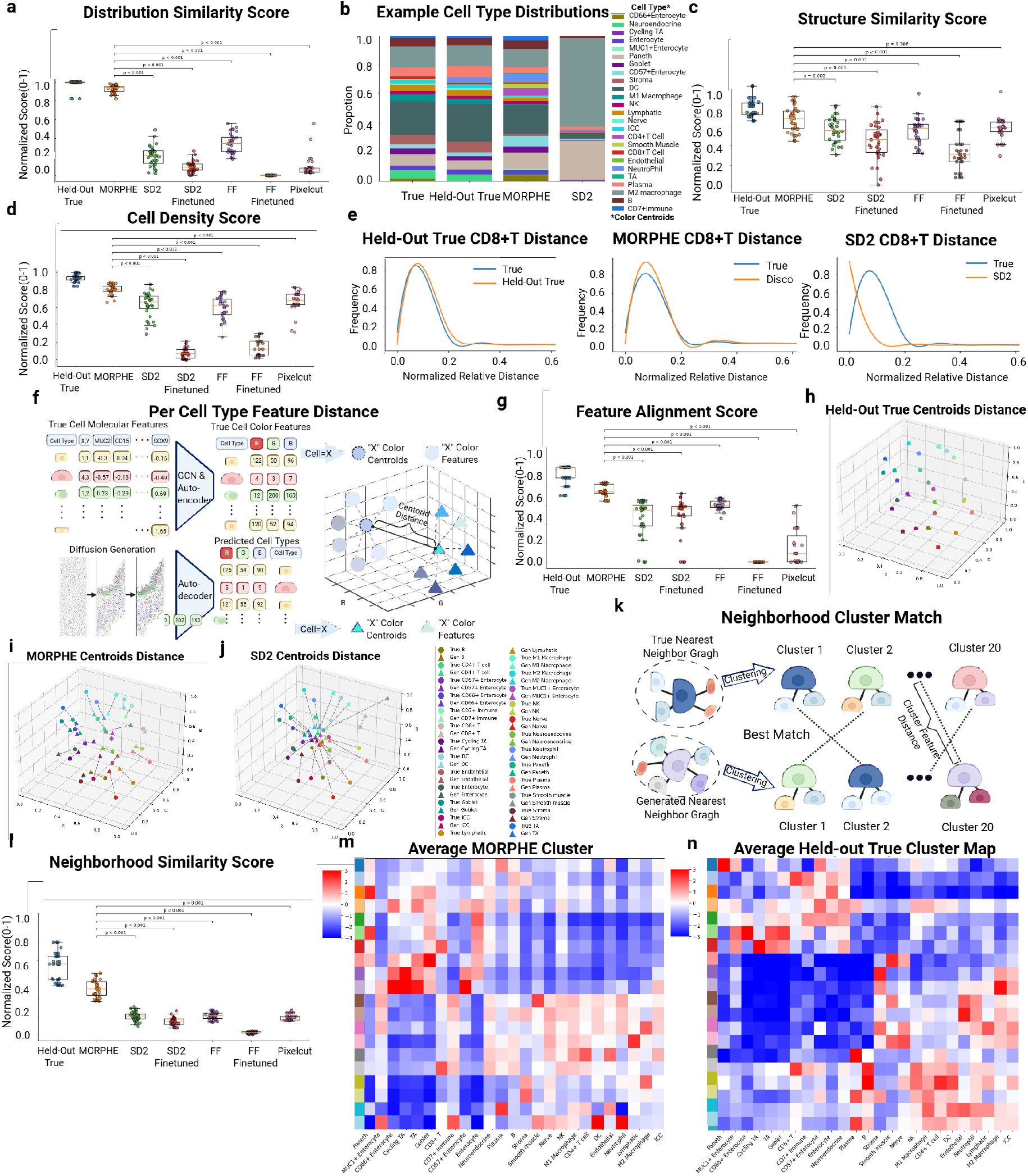
New spatial-omic generative evaluation metrics demonstrate MORPHE’s performance. **a)** Distribution Similarity Score for cell-type distributions across 31 regions (*p <* 0.001, t-test). **b)** Example of cell type proportions generated by MORPHE and SD2 compared to the held-out true and true cell types. **c)** Structure Similarity Score across 31 regions (*p* = 0.002, t-test). **d)** Cell Density Score across 31 regions (*p <* 0.001, t-test). **e)** CD8+ T spatial density distribution for the held out, MORPHE generated region, and SD2 generated region vs. ground truth. **f)** Feature Alignment Score is calculated by calculating the distance between the centroids of ground truth cell color features in the RGB space to the centroids of generated each cell type color features in the RGB space that represents embedded cell type feature consistency. **g)** Feature Alignment Score across 31 generated regions (*p <* 0.001, t-test). **h)** RGB Centroid of each cell type of the held-out true tissue, **i)** MORPHE generated tissue, and **j)** SD2 generated tissue vs. reference tissue. **k)** Neighborhood Similarity Score is calculated by measuring the distance between the shortest distance of neighborhood vector clusters between the true and generated nearest neighbor graph. **l)** Normalized Neighborhood Similarity Score across 31 generated regions (*p <* 0.001, t-test). **m)** Average neighborhood cluster heatmap for all 66 reference regions and **n)** MORPHE generated region tissue’s neighborhood in each of the clusters across the 31 regions generated. Each heatmpa show’s enrichment of cell types within each multicellular neighborhod cluster.

Second, to evaluate structural accuracy, we used a Structural Similarity Score (Learned Perceptual Image Patch Similarity (LPIPS)) [35], which is a perceptual distance measure originally developed in computer vision to quantify structural and textural similarity between images and is widely used to evaluate the fidelity of generated or reconstructed images. Here, we adapt LPIPS to assess how well MORPHE preserves spatial tissue architecture by comparing structural patterns between generated and real tissue regions. MORPHE achieved a mean score 13% higher than the SD2 (next-best alternative) (**Fig. 3c**). These results suggest that MORPHE’s sliding-window diffusion, unlike one-shot generation approaches, allows continuous directional expansion of tissues while preserving spatial coherence between old and newly generated regions.

Third, to evaluate spatial arrangement of cell populations, we used a cell-type specific density score, which computes separate distance histograms for each individual cell type, allowing us to evaluate whether the spatial positioning of each specific population is faithfully preserved [36]. The cell density scores corroborate structural accuracy of MORPHE’s generated region with the mean cell density score 0.78, and higher than the other models (**Fig. 3d**). For instance, in the CD8+ T cell population, the neighborhood distance histograms of real and MORPHE-generated tissues were more closely aligned (**Fig. 3e**), suggesting accurate spatial organization.

Fourth, to measure semantic consistency of outpainted regions, we developed the Feature Alignment Score (**Fig. 3f**), which quantifies how well cell-type-specific features are preserved. Since each cell type’s RGB values represent its classification probabilities, any diffusion-induced drift in RGB space indicates semantic distortion, such as type misclassification or overlap between classes. A consistent RGB distribution before and after generation implies that the semantic identity of each cell type is preserved. Thus we compare the Euclidean distances of the RGB centroids of each recovered cell type to that from real tissues.

Using this metric, we observe that MORPHE achieved a 15% improvement over the next best method (**Fig. 3g**), showing its ability to preserve fine-grained representations captured in molecular features of each cell type compared to other models. Indeed, by visually comparing RGB centroids across three conditions we see very little change in the held-out true tissue vs. reference tissue (**Fig. 3h**). MORPHE’s centroids remain closely aligned with the reference, maintaining type separation (**Fig. 3i**) and structure whereas SD2’s embeddings, conversely, show larger distances to held-out true RGB centroid, indicating semantic drift and loss of interpretability (**Fig. 3j**). This confirms that MORPHE preserves a more biologically meaningful and stable feature space to help preserve cell typ identities limiting semantic drift or misclassification.

Fifth, to evaluate how well spatial microenvironment relationships are preserved, we developed the Neighborhood Similarity Score. For each cell, we cluster its local neighborhood distribution (based on neighboring cell types) in both reference and generated tissue using K-Means [11]. We then match the cell neighborhood clusters using the Hungarian algorithm, which will show whether the spatial microenvironments (i.e., the local composition and organization of cell type) are preserved in the generated tissue. The score is defined by the mean centroid distance between matched clusters (**Fig. 3k**).

Using this metric, we observe that MORPHE achieved a 12% improvement over the best-performing baseline, suggesting it more accurately preserves spatial cellular neighborhoods (**Fig. 3l**).We can further appreciate the biological implications from these differences by analyzing the cell type composition within each neighborhood cluster (**Fig. 3m-n, Extended Data Fig. 5b-c**). The heatmap of MORPHE-generated tissues (**Fig. 4m**) more closely mirrors the ground truth (**Fig. 4n**), showing clear hot and cold zones that reflect biologically meaningful co-localization patterns—for example, immune-enriched regions containing macrophages and T cells, and epithelial zones enriched in enterocytes [11]. These patterns are consistent with known tissue organization and cell–cell interaction domains, indicating that MORPHE effectively captures biologically coherent spatial structures. In contrast, the SD2 output lacks such organized spatial structure, displaying mixed or diffuse clustering (**Extended Data Fig. 5c**). These results suggest that MORPHE better preserves the spatial organization and biological coherence of tissue microenvironments.

Overall, these findings demonstrate that MORPHE more faithfully preserves both the semantic identity of individual cell types and the spatial organization of tissue microenvironments compared to existing SOTA image generation models. We created two new scores that show by maintaining biologically meaningful feature embeddings and fine-grained local structure, MORPHE generates spatially coherent and interpretable tissues at single-cell resolution. This highlights its potential to bridge generative modeling with biological realism, enabling accurate and biologically grounded simulations of complex tissue architecture.

### 2.4 MORPHE Can Inpaint Arbitrarily Shaped Sub-Regions

To address the common issue of missing or damaged tissue areas in spatial omics data—often caused by sectioning artifacts, folding, detachment, fibers, or imaging errors—we developed a flexible tissue inpainting approach using MORPHE. Our method enables the restoration of arbitrarily sized missing areas, regardless of shape, by leveraging contextual information from the surrounding tissue (**Fig. 4a**).

**Fig. 4.**
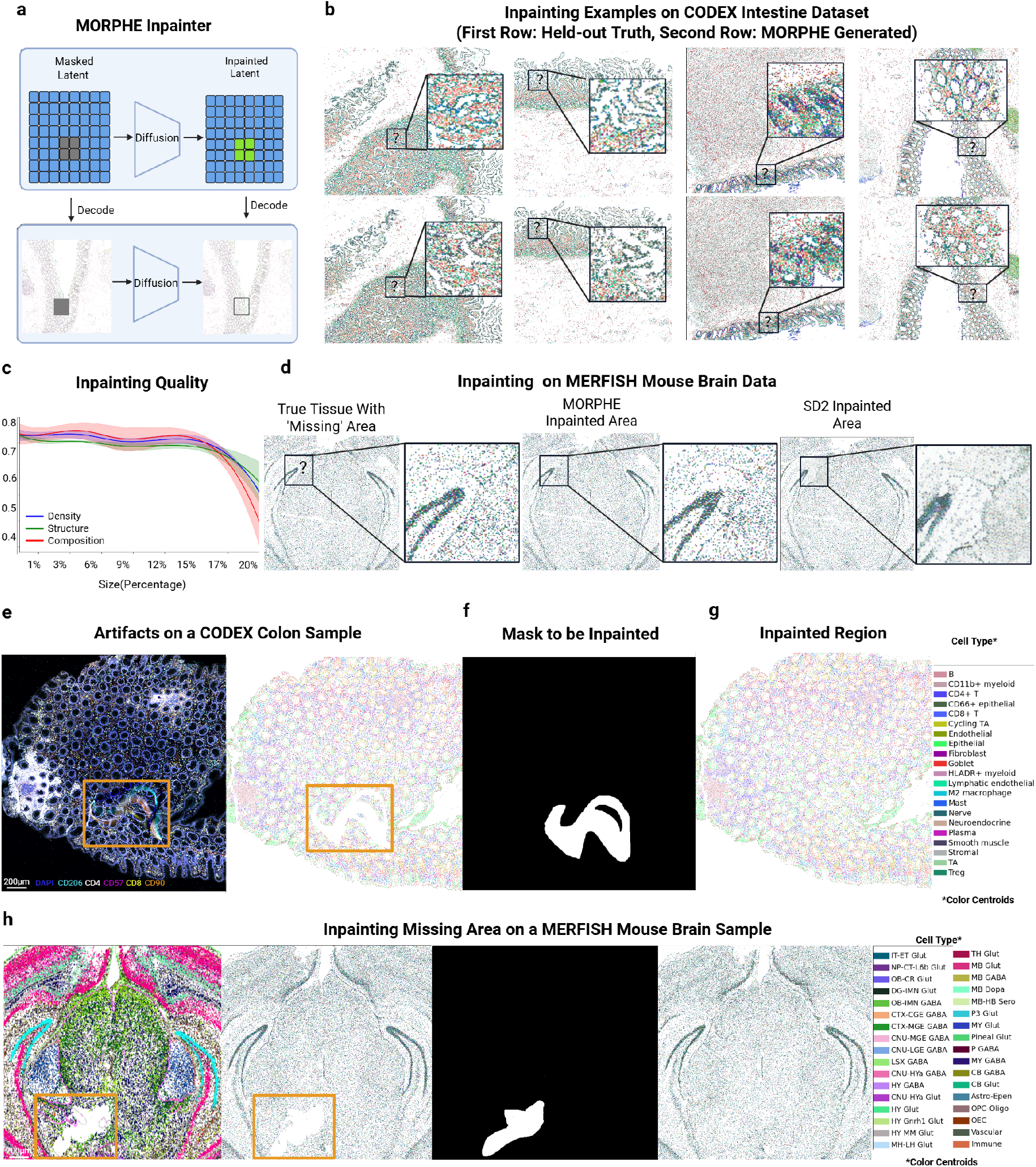
MORPHE can inpaint arbitrarily shaped sub-regions of single tissues and preserves reconstruction fidelity across multiple types of datasets. **a)** Schematic of the MORPHE inpainter: an arbitrarily shaped masked latent region is reconstructed in latent space and decoded back to pixel space. **b)** Example inpainting results on a CODEX human intestine dataset, illustrating that MORPHE restores local cellular structure and continuity; zoomed insets highlight reconstruction detail. **c)** Inpainting quality as a function of masked inpainted area (measured by cell-density similarity, structural similarity and cell-type distribution similarity). For consistency the masked regions in this analysis are square. Solid lines denote the mean across five held-out samples and shaded areas show the range (min–max) across those samples. **d)** Inpainting on the same MERFISH spatial-transcriptomics region of the mouse brain: ground truth, MORPHE inpainted result, and an inpainted result from SD2 for comparison. **e)** Example where an imaging artifact (fiber) in a CODEX healthy colon sample causes missing data: (left) original imaging with the fiber region indicated by an orange box; (right) pixelated embedding after MORPHE preprocessing (orange box). **f)**Binary mask (white) indicating the region to be inpainted. **g)** MORPHE inpainted region. **h)** Example in a MERFISH mouse-brain section with a missing region: (left) original image with the missing area boxed in orange; (second) pixelated embedding after MORPHE preprocessing (orange box); (third) binary mask (white) indicating the region to be inpainted; (right) MORPHE inpainted reconstruction.

This capability improves tissue continuity in compromised samples and enhances the completeness and downstream interpretability of spatially resolved datasets.

To evaluate the inpainting performance of MORPHE, we apply our model to the same dataset as the outpainting task which spans diverse intestinal regions and donors [11] (**Fig. 4b**). While we would not expect identical representations, MORPHE generated structurally and compositionally consistent filling of diverse areas of intestinal tissue across different cross-sections. We assessed inpainting quality using the same five metrics as we developed for the outpainting task, comparing to other methods and using a “held-out-true” tissue from a size-matched adjacent area with comparable morphology. Across all five metrics, MORPHE again consistently outperforms competing methods (**Extended Data Fig. 6a-e**).

To further understand the limits of MORPHE’s inpainting capability, we examined how the performance varies with the size of the missing region (**Fig. 4c**). When the missing region accounts for less than 17% of the total tissue area, which corresponds to approximately 8,000 valid cells on average, the model maintains stable performance across all metrics, showing little to no degradation. As the masked region increases beyond this threshold, a gradual decline is observed, particularly in Structural Similarity, which drops more noticeably—suggesting that large spatial MORPHEntinuities hinder the model’s ability to reconstruct fine structural details. Similarly, Distribution Similarity also declines with increasing mask size, indicating reduced accuracy in recapturing the original cell-type composition. These findings suggest that while MORPHE remains robust on moderate-scale inpainting tasks, restoring very large missing regions remains a more challenging problem, motivating future work on long-range structure-aware inpainting generation.

To evaluate the generalizability of the MORPHE framework beyond multiplexed spatial proteomics, we applied MORPHE to a MERFISH spatial transcriptomics dataset [9], which includes 34 annotated cell types, 58 brain sections, and over 4 million cells. With 99.69% of cells maintained post-data processing and dimension reducing, MORPHE achieved an embedding recovery accuracy of 99.05%, which means we can reconstruct the original cell type from the embedding (i.e. After embedded with GCN and Autoencoder) accurately (**Extended Data Fig. 7a**), demonstrating conservation of cell type features. In inpainting experiments on held-out sections, MORPHE produced visually sharper, more distinct, and structurally coherent cellular reconstructions (**Fig. 4d**) compared to FLUX Fill and SD2 (**Extended Data Fig. 7**).

In all preceding inpainting experiments, missing regions were synthetically created by masking sub-regions within held-out test datasets, allowing direct comparison with the corresponding ground truth. We next applied the method to an actual dataset containing genuine missing or corrupted regions arising from experimental artifacts.In a CODEX dataset of healthy human colon, a fiber introduced during imaging obstructed part of the tissue and prevented reliable cell-type annotation due to significant autofluorescence (**Fig. 4e**). However, this masked area was not rectangular as our prior examples (**Fig. 4f**). To accommodate arbitrary missing-region geometries, we represent the target region to be inpainted as a binary mask, where pixels with value 1 indicate locations to be generated and pixels with value 0 denote known prompt regions. This binary mask is provided as an additional conditioning input to MORPHE’s latent generator, enabling guided generation over irregularly shaped missing regions.

As shown in **Fig. 4g**, the inpainted region exhibits a smooth and continuous transition with the surrounding tissue, without introducing visible MORPHEntinuities or boundary artifacts while keeping the similar loop-like structure. We also demonstrated this arbitrary generation ability by inpainting a MERFISH mouse brain section (**Fig. 4h**). Again, MORPHE similarly reconstructed continuous and anatomically plausible tissue organization despite differences in imaging modality, resolution, and molecular measurement. Importantly, this example demonstrates MORPHE’s ability to handle non-ideal, non-rectangular missing areas that are not artificially defined but occur naturally in experimental data.

To quantitatively analyze the inpainting results shown in **Fig. 4g**, we adopted the same analysis strategy used for outpainting. Specifically, we computed cell neighborhood clusters for this sample (**Extended Data Fig. 6f**) and observe that the inpainted region exhibits a smooth transition (**Extended Data Fig. 6g**) and closely matches the neighborhood distribution observed in the original tissue (**Extended Data Fig. 6h**). In particular, the reconstruction preserves key crypt-associated spatial features, including the characteristic concentric arrangement of goblet-enriched, epithelial-enriched, and cycling TA–enriched neighborhoods. This is similar at a higher order biological scale where the inpainted region displays a biologically plausible transition from the middle layer toward the top layer, consistent with the organization of the surrounding tissue (**Extended Data Fig. 6i**).

To further assess the impact of inpainting on cell–cell interaction patterns, we compared pairwise distances between cells located near the artifact boundary (i.e., the edge of the missing region) with and without inpainting (**Extended Data Fig. 6k**). We observed that, in the absence of inpainting, pairwise distances are systematically increased across nearly all cell-type pairs, indicating that missing regions introduce artificial sparsity near the boundary and distort measurements of cell–cell proximity. As a representative example, the Goblet–Plasma interaction shows one of the most pronounced effects: without inpainting, some Goblet cells are forced to search farther outward to identify their nearest Plasma neighbors, artificially inflating interaction distances (**Extended Data Fig. 6l**). In contrast, after inpainting, Goblet–Plasma pairs become more densely distributed with substantially reduced distances (**Extended Data Fig. 6m**), effectively mitigating the distortion of cell–cell interaction measurements caused by missing tissue. This analysis highlights a key motivation for inpainting: to restore spatial continuity and enable more accurate quantification of cellular interactions in the presence of experimental artifacts.

Together, these results demonstrate that MORPHE’s inpainting not only reconstructs visually coherent tissue but also faithfully recovers biologically meaningful properties of the missing regions. By preserving cell-type composition, neighborhood structure, and fine-grained spatial organization, MORPHE ensures the restored areas integrate seamlessly with the surrounding tissue context and captures higher-order spatial dependencies and subtle morphological variations. As a result, the refined outputs maintain both structural continuity and biological plausibility—an essential requirement for downstream spatial-omics analyses such as quantifying cell–cell interactions and fixing microenvironmental heterogeneity where gaps or artifactual reconstructions could otherwise compromise scientific interpretation.

### 2.5 MORPHE Can Impute Continuously Between 2D and 3D Tissue Slices

In current spatial-omics imaging area extends only a few centimeters and often requires sectioning smaller sections of tissue. Moreover, most approaches are limited to 2D cross sections limiting insight to three-dimensional structure of tissues. Recently, researchers have worked around this by capturing multiple different tissue regions along 2D axis or sectioning consecutive slices and imaging for 3D representation of tissues. However, this is both time consuming and extremely costly to do. To address these limitations we designed MORPHE to be able to impute biologically plausible intermediate tissue by leveraging the structure and context of 2D or 3D anatomically adjacent regions.

For 2D imputation, unlike conventional outpainting, which expands tissue from a single prompt, we perform symmetric generation from two flanking tissue slices, progressively generating new patches inward from both ends until the gap is fully filled (**Fig. 5a**). At each generation step, MORPHE synthesizes one patch from each end, removes the oldest distal patch from the prompt, and appends the new patch for the next iteration. This bi-directional process can be iteratively applied until a predefined number of iterations is reached. In this way, MORPHE can impute long gaps and generate tissue transitions that are continuous and biologically reasonable.

**Fig. 5.**
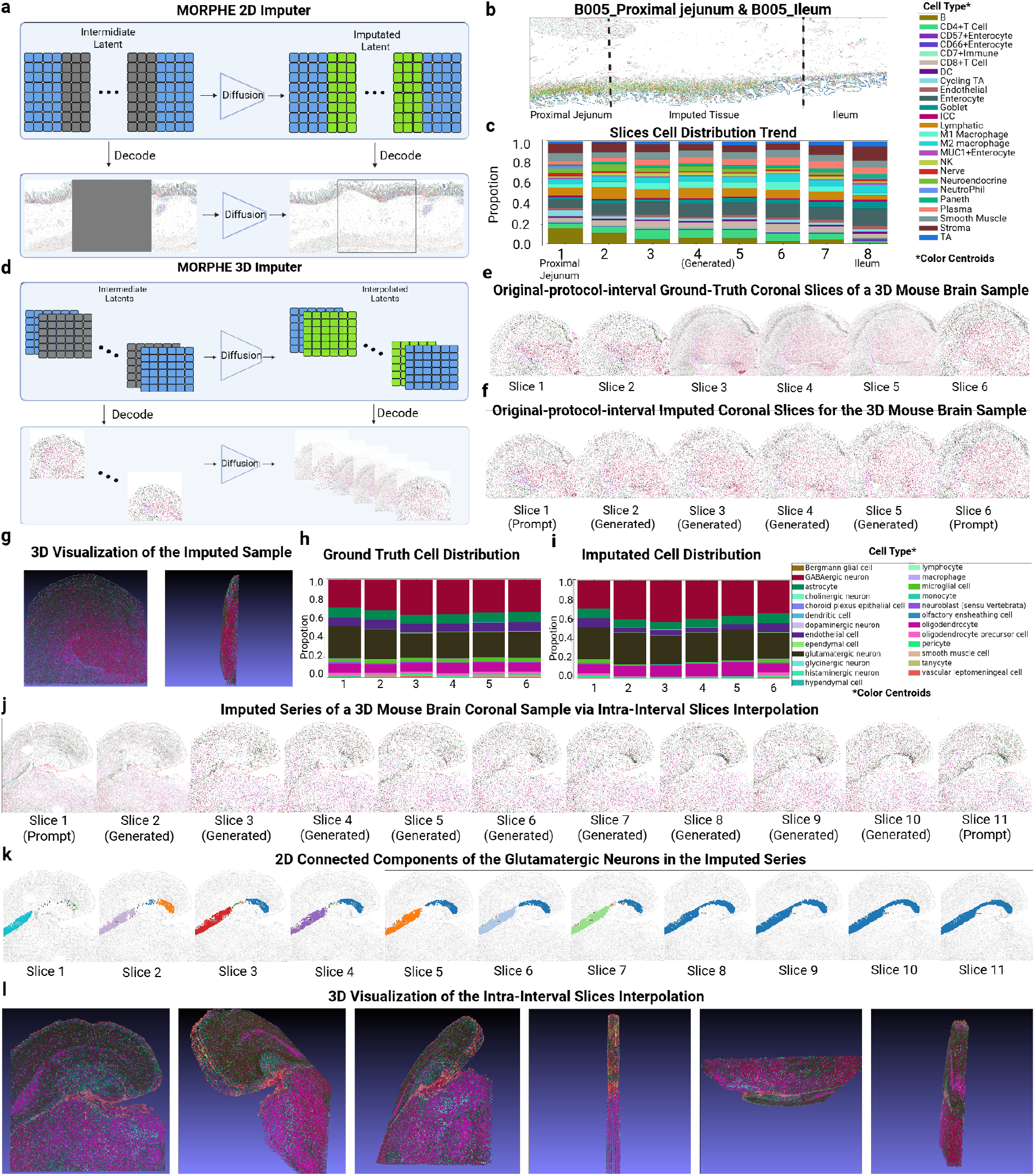
MORPHE enables continuous imputation across spatially separated tissue regions in 2D and 3D. **a)** Schematic of the MORPHE 2D imputer: a rectangle masked latent region between two endpoint prompt tissue is reconstructed iteratively in latent space and decoded back to pixel space. **b)** MORPHE imputes tissue region between Proximal Jejunum and Ileum of patient B005 in CODEX intestine dataset. **c)** Cell type distribution for individual and adjacent slices of MORPHE-imputed intestinal tissue between the two-sided prompt: B005 Proximal Jejunum and B005 Ileum. **d)** Schematic of the MORPHE 3D imputer: given two “endpoint” slices (the prompt images at either side of a missing region), MORPHE generates a sequence of intermediate slices by continuously interpolating the contribution of each endpoint. Concretely,a series of intermediate latent vectors is produced by blending the two endpoint latents with progressively changing weights (i.e., linear varying coefficients). **e)** Ground truth of an fixed-interval slices in series of a 3D MERFISH mouse brain dataset. **f)** MORPHE Imputation result of the fixed-interval slices series in **e. g)** 3D visualization of the imputation slices series in **f. h)** Cell type distribution of the ground truth slices series in **e. i)** Cell type distribution of the MORPHE imputed slices series in **f. j)** MORPHE imputation of the intra-interval slices series for another 3D MERFISH mouse brain sample. **k)** A hippocampal subregion enriched in glutamatergic neurons is shown. For each 2D slice in the series (**j**), connected components among glutamatergic neurons were computed using a 10-nearest-neighbor (10NN) criterion: two glutamatergic neurons were considered connected if one appeared within the other’s 10 nearest neighbors. Distinct connected components are indicated by different colors and number of glutamatergic neurons inside. **l)** 3D visualization of the imputation slices series in **j**.

To evaluate this capability, we applied MORPHE to multiple tissue pairs with known anatomical continuity, including the proximal jejunum–ileum pair from donor B005 (**Fig. 5b**) and the proximal jejunum–mid jejunum pair from donor B006 (**Extended Data Fig. 9a**). In the B005 proximal jejunum–ileum case, the generated intermediate region exhibited a gradual transition of tissue features between the two anatomical endpoints. This includes a reduction of the smooth muscle area where the ileum sample does not contain this.

To quantify this transition, the generated region was divided into longitudinal slices, and cell-type compositions were analyzed. CD66^+^ enterocytes were abundant in the proximal jejunum (**Fig. 5c, slice 1**) but progressively decreased across slices 2—7 and were nearly absent in the ileum (**Fig. 5c, slice 8**). Conversely, DC cells, which were sparse in the proximal jejunum but enriched in the ileum, showed a gradual increase across the intermediate slices (**Fig. 5c, slices 2—7**). Other cell types with smaller compositional differences remained largely stable. Similarly, in the B006 proximal jejunum–mid jejunum pair, plasma and DC cells exhibited a progressive increase across the generated region, while other cell types remained nearly unchanged, further supporting the smooth biological transition generated by MORPHE (**Extended Data Fig. 9b**).

For 3D imputation, unlike the iterative patch-based strategy used in 2D, we needed to enhance MORPHE to directly infer every intermediate slice between two observed endpoint slices independently (**Fig. 5d**). This is because the 3D setting provides densely sampled slice series with known ground-truth intermediates, enabling supervised learning of direct slice inference. This allows MORPHE to depart from the progressive patch-based strategy required in 2D imputation—where intermediate targets are unavailable—and instead perform independent, parallel prediction of each slice, improving stability, scalability, and reconstruction fidelity.

Given two endpoint slices, each middle slice along the axial dimension is generated independently in a single forward pass. The slice is conditioned on the two endpoints using position-aware, distance-weighted prompts that reflect its relative spatial proximity. This design provides greater flexibility and avoids error accumulation while preserving smooth and anatomically plausible transitions across slices. This is because each intermediate slice is inferred directly from the same pair of observed endpoint conditions rather than recursively conditioned on previously generated slices. As a result, prediction errors do not propagate or amplify along the depth dimension, which is a common failure mode of iterative or autoregressive strategies.

At the same time, smooth transitions are ensured by training on densely and regularly sampled slice series, where adjacent slices exhibit gradual biological variation. The model therefore learns a continuous, endpoint-conditioned manifold of valid intermediate states, allowing independently inferred slices to remain mutually consistent without explicit sequential dependence.

We applied the 3D MORPHE imputer to a coronal whole-mouse-brain MERFISH dataset comprising 183 sections and approximately 6.1 million cells from two mice. All sections from one mouse were used for training, while the entire second mouse was held out as an independent test set. In this dataset, coronal sections from a male mouse were acquired at fixed 200-*µ*m intervals. This results in a sparsely sampled 3D volume along the axial dimension where there are missing tissue sections in between sections.

To first test MORPHE’s imputation, we imputed only missing slices at the predefined data intervals (i.e., 200-*µ*m intervals), without generating any additional slices. We selected six consecutive coronal slices from a male mouse brain sample and used the first and the sixth slices as endpoint slices, which served as prompts for the 3D imputer (**Fig. 5e**, enlarged view in **Extended Data Fig. 8a**). We applied MORPHE to infer the four missing intermediate slices (**Fig. 5f**, enlarged view in **Extended Data Fig. 8b**). Visually, the imputed slices exhibit a gradual morphological transition that closely matches the ground truth. More specifically, the overall slice contour progressively expands from one endpoint to the other, and the characteristic triangular cavity in the center also increases smoothly in size across slices. We further visualized the imputed slice series in 3D (**Fig. 5g**), where the reconstructed volume shows spatial alignment with the ground truth and preserves continuous anatomical structures across the imputed region. For evaluating cell composition transition, we observed consistent shifts in cell-type composition (**Fig. 5h-i**). Major cell types, including gabaergic neuron and glutamatergic neuron, exhibit spatial distributions in the imputed slices that closely follow those of the ground truth, with comparable relative abundances across slices.

We also applied MORPHE in another case of the mouse brain 3D sections where the size of the two endpoint sections are relatively similar, but the cellular organization changes more (**Extended Data Fig. 9c**). In this case, we see that MORPHE imputation captures internal structural variation, where the intermediate slices exhibit a gradual change in internal leaf-like structures (**Extended Data Fig. 9d**). The results also show good spatial alignment and structural consistency (**Extended Data Fig. 9e**), and the overall cell-type distribution of the imputed slices closely matches the ground truth (**Extended Data Fig. 9f–g**).

To quantitatively evaluate the imputation quality, we applied the same five evaluation metrics used in previous experiments (**Extended Data Fig. 9h**). Specifically, scores for the imputed slices were computed by comparing each of the four imputed slices with its corresponding ground-truth slice. As a reference baseline, we additionally computed scores by directly comparing four intermediate ground-truth slices with the adjacent left ground-truth slice of each. Since both slices are real, this baseline reflects the natural similarity between ground-truth slices across positions, serving as a reference and sanity check for evaluating the imputed-to-ground-truth similarity. The imputed slices achieve scores consistent with levels we observed in inpainting and outpainting evaluations for each of the five metrics, closely matching the scores obtained from ground-truth-to-ground-truth comparisons (**Fig. 5h**). These results demonstrate that MORPHE can accurately recover missing tissue sections under specified intervals, producing anatomically coherent and biologically faithful 3D reconstructions.

In addition to generation of coarse grained 3D tissue reconstruction, we applied MORPHE to fully reconstruct tissues in 3D by filling in all missing sections. In the MERFISH mouse brain datase [10], the 200-*µ*m coronal interval spans multiple cell layers, while neuronal soma sizes are typically on the order of ~ 20 *µ*m [37]. Consequently, each protocol-defined interval contains approximately nine layers of cells, leaving substantial fine-scale biological variation unresolved. Thus, MORPHE needs to generate higher-density intermediate slices between two adjacent protocol-defined coronal sections, exceeding the original sampling grid.

To evaluate intra-interval imputation, we selected two adjacent coronal slices from the same male mouse brain sample (**Fig. 6j, slices 1 and 11**, enlarged view in **Extended Data Fig. 9c**) and applied MORPHE to interpolate nine intermediate slices between them (**Fig. 5j, slices 2–9**, enlarged view in **Extended Data Fig. 9c**). The generated slice series exhibits a smooth and progressive emergence of anatomical structures, including the gradual emergence of a glutamatergic neuron–enriched structure at the distal end of the hippocampus, located near the center of the reconstructed volume. Three-dimensional visualization of the interpolated slices (**Fig. 5l**) further demonstrates strong axial alignment and high structural continuity across subregions along the *z*-axis. Quantitative evaluation of cell-type composition also indicates high consistency across the interpolated slices. In particular, the cell type distribution metric achieves an average score exceeding 0.97 (**Extended Data Fig. 9i**), suggesting that intra-interval imputation preserves biologically meaningful cell-type patterns.

To quantify spatial continuity in the imputed tissue, we characterized connectivity among cells using a k-nearest-neighbor (kNN) graph constructed from their spatial coordinates. In this graph, cells are treated as nodes and edges connect each cell to its k nearest spatial neighbors, providing a local representation of tissue organization. Such kNN-based connectivity graphs are widely used to model structural continuity and transitions in biological point clouds and spatial transcriptomics data, as they capture local neighborhood relationships without imposing global shape assumptions [38, 39].Within this framework, connected components correspond to spatially coherent cellular assemblies, allowing us to assess whether specific anatomical structures remain continuous or become fragmented across slices. By analyzing the evolution of connected components across the imputed slice series, we directly evaluate whether MORPHE preserves structurally plausible spatial connectivity.

Specifically, for each 2D slice in **Fig. 5j**, we focused on the middle spoon-shaped region enriched in glutamatergic neurons at the distal hippocampal end and quantified its spatial connectivity using a 10NN-based graph representation. Connected components were then identified on each slice to characterize local spatial coherence. In the visualization, different colors denote distinct connected components derived from the 10NN graph; these colors serve only to distinguish spatially disconnected neuronal clusters and do not encode additional biological information. Across the interpolated slice series, we observe that the initially fragmented and spatially dispersed connected components in the leftmost original slice progressively decrease in number and merge across intermediate slices, ultimately forming a single dominant connected component in the reconstructed region (**Fig. 5k**). Further, we recomputed a 15NN-based three-dimensional connectivity graph on the stacked slices and visualized the resulting connected components in 3D space (**Extended Data Fig. 9j**). This analysis shows that glutamatergic neuron–enriched regions form a largely continuous and dominant 3D connected component across the reconstructed volume. This progressive consolidation of connectivity indicates that MORPHE’s intra-interval imputation yields spatially consistent neuronal organization across the interpolated volume, rather than producing disconnected or incoherent structures.

By imputing intra-interval slices, MORPHE effectively enhances axial resolution and enables finer-grained three-dimensional reconstruction beyond the original sampling density. Importantly, intra-interval imputation is performed independently for each target slice using distance-weighted conditioning on the two neighboring protocol-defined slices. This non-iterative formulation provides greater flexibility and avoids error accumulation across slices, while maintaining smooth and coherent transitions in both tissue morphology and cell-type composition.

Together, these results demonstrate that MORPHE provides a unified and flexible framework for imputing missing tissue sections in both 2D and 3D spatial omics data. By leveraging anatomical context from adjacent regions, MORPHE generates smooth and biologically reasonable transitions across spatial gaps. Importantly, MORPHE supports both protocol-aligned 3D imputation, which preserves the original experimental resolution, and intra-interval 3D imputation, which enables finer-grained reconstruction beyond the original sampling density. This flexibility makes MORPHE broadly applicable to spatial omics studies affected by sparse sampling or missing sections, facilitating more continuous analysis of tissue organization and cellular composition.

## 3 Discussion

Spatial-omics connects biological information across scale: molecules define cell state, cells communicate through local interactions, and multicellular neighborhoods assemble into higher-order tissue function. While they have substantially mapped new biology, spatial datasets are frequently incomplete maps. Cost, limited tissue availability, sectioning, and imaging artifacts yield cropped fields of view, damaged regions, discontinuous sections, and sparsely sampled 2D or 3D volumes. These gaps can distort continuity-sensitive analyses such as neighborhood calling, proximity statistics, layer quantification, and cross-sample comparisons.

To address these challenges, we developed MORPHE, a diffusion-based generative framework for spatial cellular maps. Rather than generating raw microscopy or expression values, MORPHE models the multiscale spatial organization of cellular identities derived from molecular measurements, and uses this learned prior to complete missing context. Across large spatial proteomic and transcriptomic datasets, MORPHE supports inpainting of damaged or missing regions, outpainting beyond partially imaged tissue, and interpolation across discontinuous sampling in both 2D and 3D. Together, this framework transforms fragmented measurements into continuous tissue maps that can be analyzed using standard spatial toolkits.

One practical consequence of this is that this can expand the usable fraction of archival or imperfect datasets, and reduce reliance on resource-intensive serial sectioning when the goal is to study tissue-scale organization. As spatial atlas efforts such as HuBMAP, HTAN, HCA, and SenNet continue to expand, approaches like MORPHE offer a complementary route to leverage these large datasets for resource-limited studies. For example, MORPHE can help integrate discontinuous sections and recover more continuous anatomical context from limited coverage in both 2D and 3D. Another example is the mitigation of biases introduced by experimental boundaries and artifacts. By restoring geometric continuity, MORPHE provides a unified strategy for extending and harmonizing spatial measurements across partially observed samples, enabling more comparable estimates of spatial structure and composition.

A central conceptual contribution of MORPHE is a representation bridge between discrete cell-type maps and continuous image-based generative models [26, 29]. Diffusion models operate naturally over continuous signals (e.g., RGB images, XYZ coordinates) but cannot directly operate on discrete, unordered categorical labels such as cell types. MORPHE resolves this mismatch by (i) constructing dense, pixel-resolved tissue maps that preserve cell boundaries while reducing sparsity, and (ii) learning a neighborhood- and feature-informed continuous embedding in a three-channel RGB-like space. Because this embedding is derived from probabilistic, graph-informed cell-type relationships, it shifts the generative problem from sampling arbitrary categorical labels to sampling from a continuous manifold. This latent space encodes multiscale biological information that can be decoded back to interpretable cell types and multicellular relationships.

This representation enables a second design principle: factorizing multiscale generation. MORPHE separates global tissue organization from cell-scale fidelity by combining latent-space diffusion for long-range structure with a refinement stage that restores sharp, single-cell boundaries. This cascaded design is well matched to spatial biology, where biological fidelity requires both correct composition (which identities are present) and correct geometry (how they are arranged, layered, and interfaced). Consistent with this, MORPHE better preserves boundary precision and neighborhood structure during completion than baselines that blur class boundaries or collapse toward dominant modes when operating on the same inputs.

MORPHE is to our knowledge the first approach that conditions directly on the observed spatial data to complete missing context, without requiring an external atlas for the unmeasured region. Other methods rely on predefined references, and de novo reconstruction assumes molecular similarity reflects spatial proximity—assumptions that can limit performance in heterogeneous tissues. Moreover, despite integrating a GCN, autoencoder, and cascaded diffusion stages, the framework can be trained and fine-tuned on a single NVIDIA A100 (40 GB), lowering the barrier to adoption for laboratories without extensive HPC resources. Finally, by operating on low-dimensional, spatially organized molecular representations and learned neighborhood structure, MORPHE provides a modality-agnostic approach to spatial completion that generalizes across proteomic and transcriptomic platforms.

As we enter new territory in generative spatial-omics, rigorous evaluation is especially important. Although standard quantitative metrics provide useful signals of reconstruction quality, they do not fully capture biological fidelity across scales. We therefore introduced two complementary metrics: Feature Alignment, which quantifies how well cell-type embeddings are preserved, and Neighborhood Similarity, which measures how faithfully local cellular environments are reconstructed. These measures provide a reproducible, biologically-grounded standard for assessing generative models in spatial biology and constitute a practical benchmark for future methods.

Finally, MORPHE suggests a broader recipe for generative modeling of structured spatial systems: translate discrete entities with high-dimensional features into a continuous latent representation, then learn multiscale organization with modern generative priors. While tissues motivate this work, similar representations arise in microbial biogeography, ecological surveying, public health mapping, and built-environment spatial analytics to name a few. We also applied it to public spatial Airbnb data [40] as a demonstration of this generality and expect further applications of it in the future (**Extended Data Fig. 10**).

In the future, MORPHE can be a foundation upon which additional multimodal conditioning is incorporated, including transcriptomic, proteomic, and morphological information, further enhancing integration and generation. Refinements to neighborhood modeling and diffusion scaling may improve reconstruction of large or complex regions. Furthermore, we envision MORPHE’s generative capacity to move to additional applications such as in silico perturbation studies to simulate how tissues reorganize under varying covariates such as treatment, age, or disease progression, similar to efforts ongoing in generative modeling of virtual cells [41]. The model’s ability to synthesize realistic, interpretable tissues also makes it a potential source of synthetic datasets for benchmarking and training spatial learning algorithms. In this way, MORPHE lays the groundwork for a new generation of generative models that can simulate, benchmark, and reason over spatial cell organization.

In summary, MORPHE establishes a unified generative framework for reconstructing and extending spatial-omics tissue maps at single-cell resolution. By combining neighborhood-informed probabilistic embeddings with cascaded diffusion generation and refinement, MORPHE produces completions that preserve composition, architecture, and microenvironment structure across multiple modalities and in both 2D and 3D settings. We anticipate that such generative spatial priors will become increasingly valuable as spatial datasets scale in size and diversity, serving as computational “connective tissue” that helps researchers correct artifacts, harmonize incomplete measurements, and explore how local cellular rules give rise to emergent tissue-scale organization.

## 4 Methods

### 4.1 Overview

In the MORPHE pipeline, tissue coordinates are first standardized across tissue regions through scaling and padding, producing square spatial layouts compatible with image-diffusion backbones. On this foundation, a graph convolutional network (GCN) [42] models each cell as a node in a spatial graph, learning classification probabilities that jointly capture biomarker features and local neighborhoods. These probabilities are further transformed by an autoencoder into a compact three-dimensional latent space, where each dimension corresponds to the R/G/B channel, forming an interpretable image representation of the tissue. Building on this representation, a cascaded diffusion model [26, 29] performs spatial-aware generation to reconstruct missing regions, yielding tissue maps that are spatially continuous and biologically coherent.

### 4.2 Data preparation

Spatial omics data describe each cell by its centroid coordinates (*x, y*), annotated cell type, and a vector of protein/mRNA-biomarker intensities. To enable image-based modeling, each tissue region was projected onto an M×M grid, where M corresponds to the side length of the largest square that closely matches the upper coordinate bounds of the original dataset (In CODEX Intestine Dataset: M=9000; In MERFISH Mouse Brain Dataset: M=3000). In this grid, each pixel represents either a cell or an empty space, allowing spatial structures to be visualized as images.

For tissue regions smaller than M×M, we padded the grid by adding empty pixels—positions without any corresponding cells—to reach the target dimensions while maintaining the correct spatial alignment of existing cells.

For regions larger than M×M, we cropped the data to fit the standardized grid. Cropping was performed selectively to minimize cell loss, focusing on subregions that contained the densest and most biologically informative portions of the tissue.

This adaptive padding-and-cropping procedure ensured that all samples shared a consistent spatial format without distorting their intrinsic geometry, enabling reliable image-based modeling across diverse tissue regions. The grid was first downsampled to 1024 × 1024 using linear scaling:

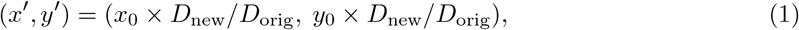

where *D*_orig_ = 9000 and *D*_new_ = 1024. When multiple cells were mapped to the same downsampled coordinate, we first computed the precise low-resolution position corresponding to each cell. In cases of conflicts—where multiple cells fell into the same grid—we retained only the cell whose computed position was closest to the center of that pixel, discarding all others. To meet diffusion model input and GPU memory limits, the 1024×1024 maps were further reduced to 512×512 using nearest-neighbor interpolation. For conflicting cells, we assign each cell to the nearest available empty pixel.

### 4.3 Graph convolutional network (GCN) for Classification Probabilities

To capture spatial and molecular relationships between cells, MORPHE models each spatially resovled cell map as a graph, where nodes represent individual cells and edges connect spatially proximate neighbors. Each node is associated with cell type, coordinates, and a feature vector of protein/mRNA-biomarker intensities, and the graph is constructed using a *k*-nearest-neighbor (KNN) [43] algorithm based on Euclidean distance.

The graph convolutional network (GCN) [42] comprises four layers: a fully connected input layer, two graph convolutional layers for message passing among neighboring cells, and a linear classifier projecting features to *K* cell-type logits. The graph convolution at iteration *l* is defined as:

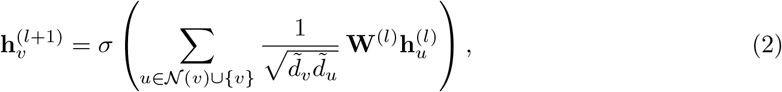

where 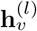 is the feature vector of node *v* at iteration *l*, 𝒩 (*v*) denotes the set of neighbors of (excluding *v* itself), 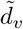 and 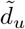 are degrees computed on the adjacency with added self-loops (i.e. *Ã* = *A* + *I*), **W**^(*l*)^ is a learnable weight matrix, and *σ*(·) is a ReLU activation.

The network is trained using cross-entropy loss:

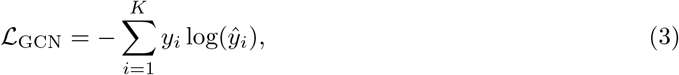

where *y*_*i*_ and *ŷ* _*i*_ denote true and predicted classification probabilities. Using four layers, a hidden dimension of 1024, and *k* = 20, the GCN achieved the best classification performance based on this cross-entropy loss.

### 4.4 Autoencoder for embedding

The autoencoder module learns a compact three-dimensional representation of the GCN-derived cell-type probability vectors. Given an input vector **p** ∈ ℝ ^*N*^, where each element corresponds to the probability of a specific cell type and *N* is the number of cell types, the encoder *E*_AE_ projects **p** into a latent embedding **z** ∈ ℝ^3^, and the decoder *D*_AE_ reconstructs the original distribution 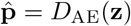.

Both the encoder and decoder are implemented as four-layer fully connected networks equipped with Layer Normalization and ReLU activations after each linear transformation. The encoder follows an expansion–compression architecture, progressively increasing the hidden dimension to 512 before symmetrically contracting toward a three-dimensional bottleneck, while the decoder mirrors this process in reverse to restore the original dimensionality. This symmetric configuration facilitates smooth information flow and stabilizes optimization during training.

To shape the three-dimensional latent manifold, we explicitly separate the cluster regularizer into intra-class cohesion and biologically weighted inter-class repulsion. The cluster regularization is defined as

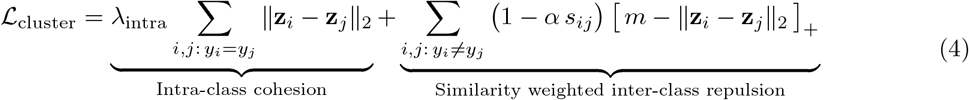

where ℒ_cluster_ represents the overall cluster regularization term. The first component, weighted by *λ*_intra_, minimizes the Euclidean distance ∥**z**_*i*_ − **z**_*j*_∥_2_ between embeddings of the same cell type (*y*_*i*_ ≠*y*_*j*_), promoting intra-class compactness. To prevent over-clustering, the penalty is intentionally set to be mild (e.g., *λ*_intra_ = 0.1), ensuring that embeddings of the same cell type remain close but not collapsed to a single point. The second component, *Biologically weighted inter-class repulsion*, encourages embeddings of different cell types (*y*_*i*_ ≠ *y*_*j*_) to maintain spatial separation in the RGB manifold through a margin-based constraint. However, this repulsive force is adaptively relaxed according to biological similarity, where *α* scales (e.g., *α* = 0.1) the penalty based on *s*_*ij*_ = cos(**p**_*i*_, **p**_*j*_)—the cosine similarity between their probability distributions. This design ensures that while distinct cell types are kept apart, those sharing similar probabilistic (indicating similar biomarkers) remain relatively closer, preserving the global biological semantic structure in latent space.

The full autoencoder objective combines the reconstruction fidelity, measured by the Kull-back–Leibler divergence [32], with the cluster regularizer:

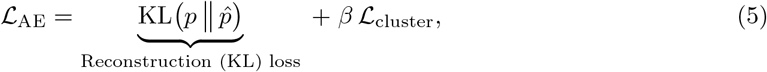

where *β* balances the influence of the clustering constraint against the reconstruction objective. This formulation ensures faithful probability reconstruction while structuring the latent manifold such that cells of the same type cluster tightly, biologically similar types are positioned nearby, and dissimilar types are well separated in the three-dimensional RGB embedding space.

Finally, the resulting latent vector *emb* = (*emb*_*R*_, *emb*_*G*_, *emb*_*B*_) is linearly normalized to [0, 1] and denormalized to RGB channels [0, 255], producing visually interpretable tissue maps used by the downstream diffusion reconstruction pipeline.

### 4.5 Cascaded diffusion process

To address the reconstruction of missing regions in cell maps, we reformulate the problem as an embedded image generation task and propose a cascaded diffusion pipeline designed to synthesize biologically consistent regions. Each cell map is treated as an image, and the model learns to generate masked regions based on visible spatial context.

#### 4.5.1 Inputs and masking

Let *I* ∈ ℝ ^*C*×*H*×*W*^ denote the input cell-map image and *M*_pix_ ∈ {0, 1}^*H*×*W*^ the binary pixel mask, where *M*_pix_(*x, y*) = 1 for masked pixels and 0 otherwise. The masked input is defined as:

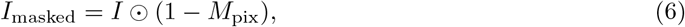

where ⊙ denotes elementwise multiplication. Each mask is parameterized by its aspect ratio *r*, area *a*, and coordinates (*x*_1_, *y*_1_, *x*_2_, *y*_2_).

#### 4.5.2 VAE Encoding and Latent Mask

A pretrained VAE encoder *E*(·) [26, 44] maps the image (3, 512, 512) into latent space (4, 64, 64):

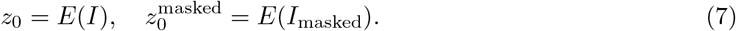

The pixel-space mask is converted to a latent-space mask *M* = *D*(*M*_pix_) using a deterministic downsampling operator *D*(·), so that 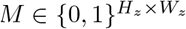 aligns with the latent resolution.

#### 4.5.3 Spatial Prompt Construction

The spatial feature encoder *S*(·) extracts contextual information from the masked input image *I*_masked_. The encoder is implemented as a residual convolutional neural network [45] that progressively expands the feature dimension from 64 to 736 channels while reducing the spatial resolution from 64 ×64 to 8 ×8. The resulting feature maps are flattened into a sequence of 64 tokens, each enriched with a two-dimensional positional encoding and normalized using Layer Normalization to stabilize training and preserve spatial correspondence.

The extracted image features are represented as

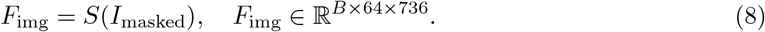

A single layer perceptron *E*_coords_(coords) encodes the geometric parameters of the masked region—as an encoded binary mask—into a compact representation:

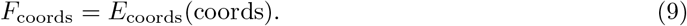

The two feature sets are concatenated along the feature dimension to form the spatial prompt:

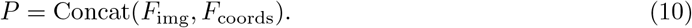

This spatial prompt provides the conditional UNet with both visual and positional context, enabling precise localization and reconstruction of masked regions.

#### 4.5.4 Latent Diffusion

Following the DDPM formulation [29], we first operate in the latent space. Given a masked latent representation 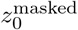, the forward process with Gaussian noise *ϵ* ~ 𝒩 (0, *I*) is defined as:

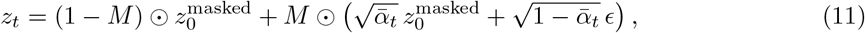

where *M* ∈ {0, 1} ^*H*×*W*^ denotes the mask. A conditional UNet *U*_1_(.) adapted from Stable Diffusion v1.5 [26] predicts the clean latent:

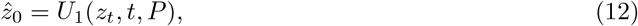

where *P* is the spatial prompt. The training objective is:

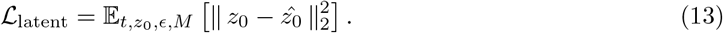

#### 4.5.5 Diffusion-Based Pixel Decoder

After the latent diffusion stage produces a reconstructed latent tensor 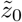 of size 4×64×64, this latent serves as the conditioning signal for the pixel-domain diffusion model.

To embed the latent with position information, a four-channel sinusoidal positional encoding [46] is defined as

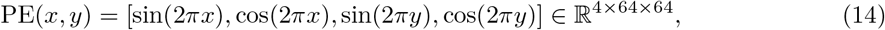

and is concatenated with 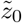 to form an eight-channel input *z*_in_ ∈ ℝ^8×64×64^.

To capture both fine and coarse spatial context at the 64×64 latent resolution before constructing the conditioning pyramid, a dilated residual block [47] refines this representation. The block normalizes and activates *z*_in_, projects it through a 1×1 convolution, and processes the resulting intermediate feature map with three parallel 3 × 3 convolutions with dilation rates 1, 2, and 4. The outputs of these branches are concatenated, normalized, re-activated, and fused by a final 1 × 1 convolution to yield

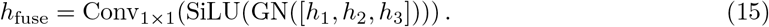

A residual update then produces the refined latent representation *x*_64_ = *z*_in_ + *h*_fuse_ ∈ ℝ^*C*×64×64^, where *C* is the internal feature dimension.

To construct multi-scale conditioning features, *x*_64_ is projected by a 1×1 convolution into 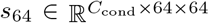, where *C*_cond_ is the conditioning-channel width. The tensor is then upsampled sequentially to 128×128, 256×256, and 512×512 resolutions using bilinear interpolation, and at each step refined by a 3×3 convolution followed by a residual block identical in structure to the 64×64 block. These intermediate representations are projected through 1×1 convolutions to obtain *s*_128_, *s*_256_, and *s*_512_, which together with *s*_64_ form the conditioning set cond feats ={ *s*_64_, *s*_128_, *s*_256_, *s*_512_}as the dominant prompt for a conditioning UNet.

The conditioning UNet *U*_2_ reconstructs the clean image 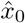 using (1) the noisy input *x*_*t*_ of size 3×512×512, (2) a sinusoidal timestep embedding 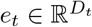 following diffusion models [29], and (3) the conditioning set cond feats. The embedding is passed through an MLP and injected into all residual blocks.

The encoder of *U*_2_ progressively reduces spatial resolution from 512×512 to 64×64, applying residual blocks at each scale and passing skip-connections to the decoder. At the 256×256 and 128×128 layers, the conditioning tensors *s*_256_ and *s*_128_ are concatenated before the residual blocks to provide scale-appropriate spatial context. At the 64×64 bottleneck, a multi-head attention module [48] normalizes the feature map, reshapes it into a token sequence, applies self-attention, and restores it to a 64×64 grid before concatenating *s*_64_ and applying two additional residual blocks.

The decoder mirrors the encoder by upsampling the representation to 128×128, 256×256, and 512×512 resolutions, again applying residual blocks per scale. At each resolution, the decoder concatenates the upsampled representation with the skip-connection and the corresponding conditioning tensors (*s*_128_, *s*_256_, and *s*_512_). A final 3×3 convolution followed by a tanh activation yields the reconstructed RGB output 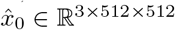.

Training follows the DDPM denoising objective [29]. For each timestep, a noisy image is formed as

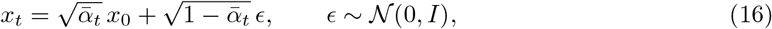

and the pixel-space decoder is trained using the squared-error loss

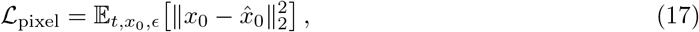

ensuring that the reconstruction remains consistent with both the latent-stage output and the multi-scale conditioning.

### 4.6 Inpainting Inference

Inpainting restores missing area in the single tissue according to surrounding context.

Given an input image *I* ∈ ℝ^*C*×*H*×*W*^ and a binary mask *M*_pix_ ∈ {0, 1}^*H*×*W*^, the visible region is

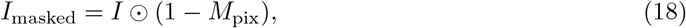

where ⊙ denotes elementwise multiplication. The VAE encoder *E*(·) projects both the full and masked images into latent space, producing

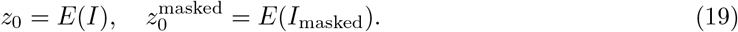

The pixel-space mask is downsampled to latent resolution using a deterministic operator *D*(·) to obtain *M* = *D*(*M*_pix_). At inference, the diffusion model begins from Gaussian noise *z*_*T*_ ~ 𝒩 (0, *I*) and iteratively denoises the masked latent using DDPMSchedulerStep [29] through *U*_1_(·):

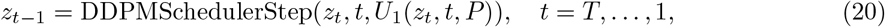

where *P* = Concat(*F*_img_, *F*_coords_) is the spatial prompt constructed in Section 4.5.3. After the reverse process, the reconstructed latent 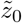 represents a complete cell map in latent space. The latent output 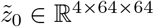is passed through a lightweight latent adapter to obtain multiscale conditioning features

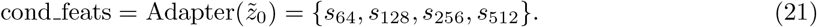

Starting from Gaussian noise *x*_*T*_ ~ 𝒩 (0, *I*), the pixel-level diffusion decoder *U*_2_(·) performs iterative denoising using DDPMSchedulerStep [29] conditioned on these features:

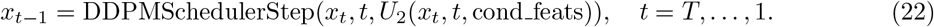

After all timesteps, the final clean image 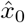 constitutes the inpainted reconstruction.

### 4.7 Outpainting Inference

Outpainting extends spatially resolved tissue maps beyond their original field of view by iteratively masking and expanding latent representations within the cascaded diffusion framework.

The input image *I* ∈ ℝ^*C*×*H*×*W*^ serves as the initial visible region, and a normalized bounding box (0, *y*_1_, 1, *y*_2_) or (*x*_1_, 0, *x*_2_, 1) defines the border area to be generated, depending on the expansion direction. Empirically, generating approximately 5% of the total area per iteration yields the most stable outpainting performance.

This bounding box is converted into a latent-space binary mask 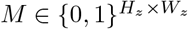, and the masked latent is initialized as *z*_0_ ⊙ (1 −*M*), preserving the original content while leaving the border region empty.

At each iteration, the masked region is reconstructed in latent space, and the newly generated patch is concatenated to the visible region along the selected expansion direction.

To maintain a fixed latent width of 64, the farthest portion of the prompt corresponding to the oldest context is simultaneously removed during concatenation, ensuring a constant input size.

This sliding-window mechanism allows progressive spatial expansion while keeping the model’s receptive field consistent across iterations. The iterative masking and concatenation process continues until the desired extension length is reached.

The generation process of each iteration follows the same cascaded denoising procedure as in inpainting [29].

### 4.8 2D Imputation Inference

2D imputer reconstructs missing intermediate tissue between two spatially separated regions by iteratively masking and stitching latent representations within the cascaded diffusion framework.

The input image *I* ∈ ℝ^*C*×*H*×*W*^ is composed of two adjacent tissue slices—*I*_*L*_ and *I*_*R*_—concatenated along one spatial axis, with an empty gap deliberately left between them.

The combined image is encoded into latent space, producing *z*_0_ = *E*(*I*). A normalized bounding box (*x*_1_, 0, *x*_2_, 1) defines the central gap corresponding to the missing region, which is converted into a latent-space binary mask 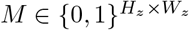.

The masked latent is initialized as *z*_0_ ⊙ (1 − *M*), preserving the visible tissue on both sides while leaving the central area empty.

At each iteration, the masked gap region is reconstructed in latent space, and the newly generated patch is stitched between the left and right latent contexts.

To maintain a fixed latent width of 64 (the shape of is fixed 4×64×64), the outermost columns of the latent tensor are cropped after each update, and a zero-filled latent gap is reintroduced as the next mask.

This iterative masking and stitching process continues until the gap between *I*_*L*_ and *I*_*R*_ is completely filled. The generation process of each iteration also follows the same cascaded denoising procedure as in inpainting [29].

### 4.9 3D Imputation Inference

The 3D imputer infers intermediate slices by interpolating latent representations between two known endpoint slices. Given a sparsely sampled 3D series, the 3D imputer reconstructs missing intermediate slices within a volumetric sample by leveraging prompt-weighted conditioning to generate slices at different spatial locations along the third dimension.

Let the observed slice series be indexed along the axial dimension, where two known endpoint slices at positions *z*_*i*_ and *z*_*j*_ bound a set of missing intermediate positions {*z*_*t*_}_*i<t<j*_. Each observed slice is encoded into latent space, yielding a set of latent representations {*z*_*k*_}.

For a target missing slice at position *z*_*t*_, the imputer conditions generation on both endpoint prompts using distance-aware weighting. Specifically, the relative distances between the target position and the two endpoints are computed as:

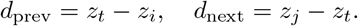

The contribution of each endpoint prompt is then determined by normalized weights inversely proportional to these distances:

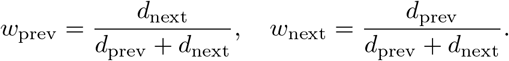

Using these weights, the latent prompts corresponding to the two endpoint slices are linearly combined to form a position-aware conditioning signal for slice *z*_*t*_. Slices closer to the left endpoint are thus more strongly influenced by the left prompt, while slices closer to the right endpoint receive higher contribution from the right prompt.

Conditioned on this weighted prompt, the imputer generates the latent representation of the missing slice following the same cascaded denoising procedure as used in inpainting [29]. This process is repeated independently for each missing slice, resulting in a continuous and spatially coherent reconstruction of the full 3D slice series.

### 4.10 Application to Airbnb Listing Dataset

To evaluate the generalizability of MORPHE beyond spatial omics data, we applied the framework to a publicly available Airbnb listing map dataset [40]. The dataset comprises two-dimensional maps from 116 cities, where each point corresponds to a rental property annotated with multiple attributes, including price, geographic location, host characteristics, number of rooms, and rental type.

To construct discrete classification identities compatible with the MORPHE pipeline, we performed unsupervised clustering on the listing feature space and partitioned properties into ten categories (Extended Data Fig. 10a). These clusters were treated as categorical types analogous to cell-type labels in spatial omics data. For each city-level region, we generated a two-dimensional atlas in which each spatial coordinate was assigned a cluster identity. The categorical labels were then embedded into the MORPHE RGB-like three-channel representation following the same encoding procedure used for biological datasets (Extended Data Fig. 10b).

We first evaluated model performance using an inpainting task. For each of 16 selected regions, a spatial subregion was masked and reconstructed using the trained generative model (Extended Data Fig. 10c). Reconstruction quality was assessed by comparing structural organization and spatial distribution patterns between inpainted areas and their corresponding reference regions (Extended Data Fig. 10d).

To further assess the framework in a scenario involving incomplete spatial coverage, we applied MORPHE to the Toronto listing map (Extended Data Fig. 10e). A region lacking data in the upper-left portion of the map was selected for outpainting. The surrounding areas exhibited a regular grid-like spatial structure, providing contextual constraints for generation. The model was conditioned on the observed neighborhood and used to generate the missing region (Extended Data Fig. 10f–g).

### 4.11 Metrics

### Structural Similarity Score

To quantify perceptual structural similarity between the generated image *I*_gen_ ∈ ℝ^3×*H*×*W*^ and the ground-truth image *I*_true_ ∈ ℝ^3×*H*×*W*^, we adopt the Learned Perceptual Image Patch Similarity (LPIPS) metric [49]. LPIPS measures distances in deep feature space extracted by a pretrained network (i.e., VGG [50]), providing a perceptually aligned metric superior to pixel-level measures such as SSIM [51] or PSNR:

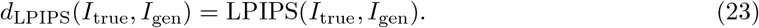

Here, *d*_LPIPS_ denotes the perceptual distance between two RGB images computed by the LPIPS network. Since smaller LPIPS values indicate higher similarity, we define the structural similarity score as

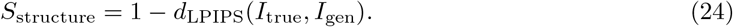

*S*_structure_ ∈ [0, 1] provides a normalized measure of perceptual similarity, with larger values indicating higher structural fidelity.

### Cell Density Consistency Score

To assess whether the generated spatial maps preserve the local neighborhood structure of cells, we compute, for each cell type *t* ∈ { 1, …, *T*}, the mean distance to its *k* nearest neighbors in both real and generated maps, following the density-based neighborhood modeling concept used in Nicheformer [36]. For each type, we construct normalized histograms 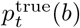 and 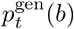 (with bin width Δ) of the mean neighbor distances. The Kullback–Leibler (KL) divergence between them quantifies the discrepancy:

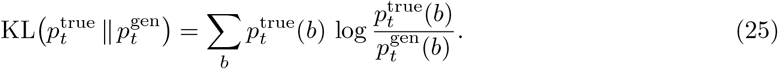

Here, 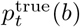 and 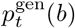 denote the normalized frequency of distances within bin *b* for cell type *t* in the true and generated maps, respectively. A per-type similarity score is then defined as

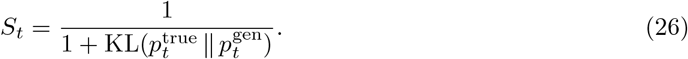

*S*_*t*_ provides a bounded similarity score in [0, 1], with higher values corresponding to more similar local density distributions.

If a given cell type is absent in the generated map, we apply a penalty term 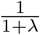 with *λ >* 0 to discourage missing types, consistent with the implementation:

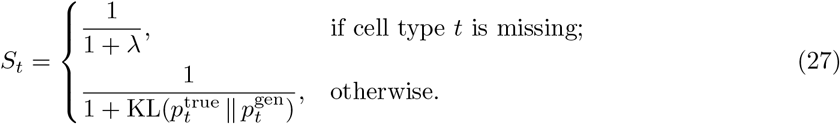

Here, *λ* is a penalty hyperparameter controlling the penalty strength for missing types. The final cell density consistency score combines per-type and global distance distributions as

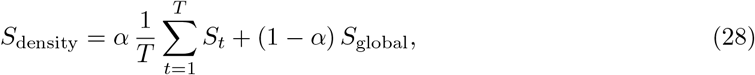

where *S*_global_ measures the similarity of global (all-cell) distance distributions, and *α* ∈ [0, 1] balances type-specific and global contributions.

### Cell Type Distribution Similarity Score

Global cell type proportions are compared between true and generated maps by KL Divergence or JSD. This KL/JSD-based comparison is commonly applied to quantify compositional differences in cell-type distributions [32, 52–54].

To handle missing cell types robustly, we introduce a penalty on absent types in the generated distribution.

Let **p** = (*p*_1_, …, *p*_*T*_) and **q** = (*q*_1_, …, *q*_*T*_) denote the normalized cell type histograms for true and generated maps, with small *ϵ* (i.e. 1e-8) added for numerical stability. We compute the penalized KL divergence as

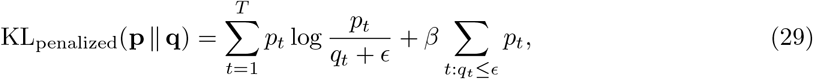

where *β* is a penalty weight that increases when a cell type present in the ground truth is missing from the generated map. Finally, the composition similarity score is defined as

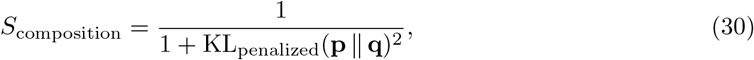

so that higher values indicate greater similarity. Here, *S*_composition_ ∈ [0, 1] and the squared denominator amplifies penalties for larger divergences.

### Feature Alignment Score

To evaluate visual consistency between generated and true spatial maps in color space, we propose a Feature Alignment Score that measures the discrepancy between the mean RGB values of each cell type. For every cell type *t* ∈ { 1, …, *T*}, we compute its centroid in RGB space from both the ground-truth image *I*_true_ and the generated image *I*_gen_ as 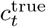 and 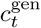 :

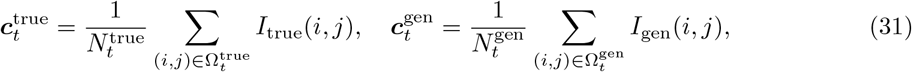

where Ω_*t*_ denotes the set of pixel coordinates assigned to cell type *t*, and *N*_*t*_ is the number of such pixels. The Euclidean distance between corresponding centroids quantifies the color deviation:

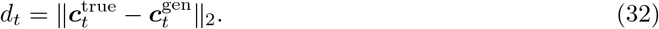

If a cell type is absent from either image (i.e., 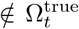 or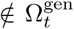), a fixed penalty value *λ* is assigned to its distance:

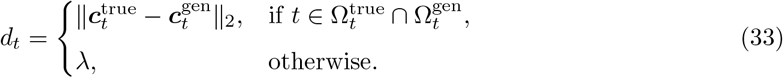

This penalty ensures that missing or misrepresented cell types contribute a constant penalty to the overall discrepancy, preventing overly optimistic scores in cases where rare types are lost during generation. In our implementation, *λ* is empirically set to 2.5, corresponding to a perceptually moderate RGB deviation.

The overall score is converted into a bounded similarity measure using a linear mapping function *ϕ*(·):

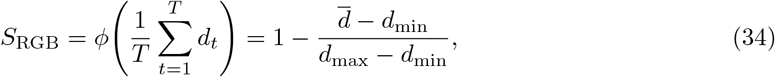

where 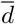 is the mean centroid distance across all cell types, and *d*_min_ and *d*_max_ define normalization bounds. The mapping ensures *S*_RGB_ ∈ [0, 1], with higher values indicating stronger alignment between generated and ground-truth color features. This metric captures per-type chromatic fidelity in an interpretable manner, linking image-space color coherence to biological structure preservation.

### Neighborhood Similarity Score

We further introduce the Neighborhood Similarity Score to assess whether the generated tissue reproduces realistic higher-order cellular co-localization patterns.

For each map, we extract the local neighborhood composition of each cell by computing the histogram of neighboring cell types within a fixed radius *k*:

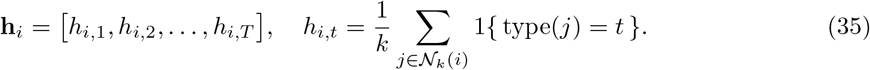

Here, 𝒩_*k*_(*i*) denotes the set of the *k* nearest neighbors of cell *i* and *h*_*i,t*_ represents the local proportion of cell type *t*. These histograms are then clustered using MiniBatch *k*-means to form *C* distinct *niche clusters* representing characteristic microenvironmental compositions. For the true and generated maps, we obtain cluster centroids 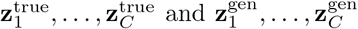. The optimal cluster correspondence is determined by solving a linear sum assignment (Hungarian) problem [55] that minimizes the total centroid distance:

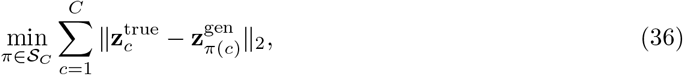

where *π* is a permutation of {1, …, *C*} defining the cluster matching. Let *D*_total_ denote the minimum summed distance. We then define the similarity score as

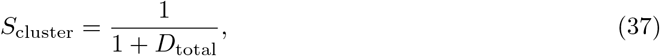

where a larger score indicates better correspondence between the compositional cluster structures of true and generated tissues. This metric captures high-level topological consistency of local microenvironments and is particularly useful for benchmarking generative spatial models that aim to reproduce tissue-level organization of heterogeneous cell types.

## 5 Data availability

The original CODEX human intestine datasets presented in this study can be found in the online repository at Dryad https://doi.org/10.5061/dryad.pk0p2ngrf

MERFISH mouse datasets are from Ref [9].

3D whole mouse brain datasets are from Ref [10].

## 6 Code availability

MORPHE was written in Python v.3.9 using the PyTorch library. The source code including finetuning Stable Diffusion 2 and Flux.1 Fill dev is available on Github at https://github.com/HickeyLab/MORPHE. All models in this work are trained on 1 A100 GPU with 40GB display memory.

## 7 Acknowledgments

This work was supported by the US National Institutes of Health (grant nos. 3U54AG07593, 3OT2OD033759-01S4, 3OT2OD033759-01S5, 1U01-AI186999-01, 1U54-AI191253-01); the National Science Foundation CAREER Award (grant no. 2440733); the V Foundation (V2025-019); the Human Frontier Science Program Early Career Grant (RGEC26/2025-); and start-up support from Duke University Department of Biomedical Engineering for JWH. M.B. gratefully acknowledges the support of the Swiss National Science Foundation (SNSF) starting grant TMSGI2 226252/1 and the SNSF grant IC00I0 231922. M.B. is a CIFAR Fellow in the Multiscale Human Program.

