## Supplemental Information for "MORPHE: Bridging Image Generation and Spatial Omics for Tissue Synthesis"

In the format provided by the authors and unedited

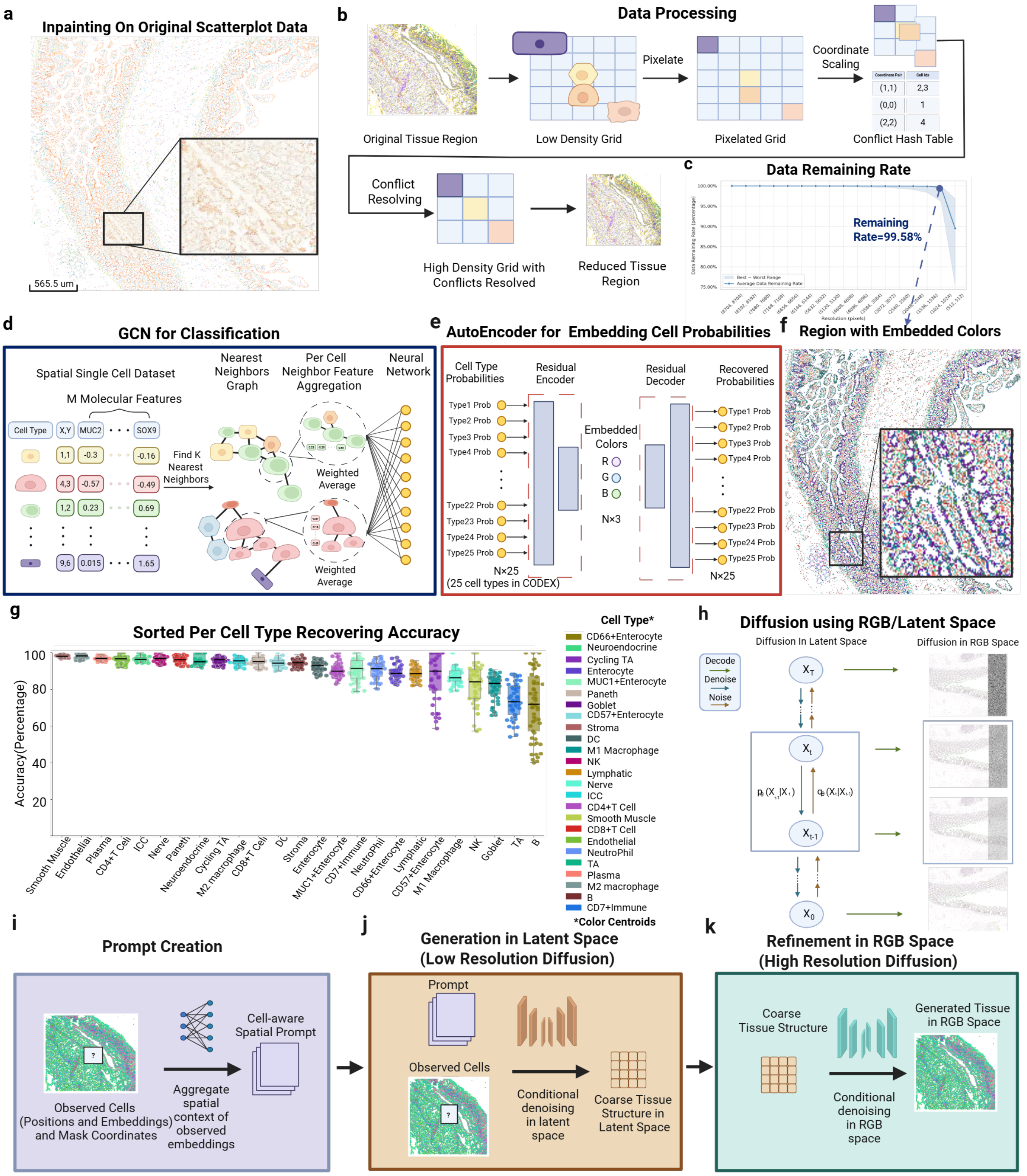


**Extended Data Fig 1** Visualization of the CODEX Intestine dataset and similarity check across dataset for validating MORPHE's generalizability. **a)** Stable Diffusion 2 inpainting output performed on raw plotted spatial proteomics data. **b)** MORPHE performs pixelation and resolution reduction by starting with cell centroid positions in a low resolution grid, representing each cell by a pixel at its centroid location, mapping these pixels with coordinate scaling to a lower dimensional grid, and resolving collisions at lower dimensional coordinates with distance priority, removing the cells which map to a position but do not have the highest proximity. **c)** Resolution reduction across multiple scales for the CODEX intestine dataset from its original resolution. At 1024x1024, MORPHE preserves 99.58% of cells. **d)** A neural network that utilizes a cell's k nearest x,y coordinate neighbors aggregates the clusters' molecular markers to classify the cell's cell type generating cell type probabilities in the last layer of the network. **e)** An autoencoder takes the last layer of the GCN pseudo probabilities, N yellow dots representing a cell's N (N=25 for intestine) different cell type probabilities, and reduces them to dimension 3, represented in embedded colors RGB. The decoder reverses this process taking the 3 dimensions back to N (N=25 for CODEX intestine dataset) dimension cell type probabilities. **f)** An intestinal region colored by using the RGB embedded cell type colors using this GCN approach. **g)** Sorted accuracy in GCN cell type prediction for each of the cell types in the CODEX intestine dataset with data points for each of the 66 tissue regions. **h)** MORPHE trains a latent diffusion process beginning from noise-corrupted embeddings. During training, noise is iteratively added to clean embeddings indexed by diffusion time t; during generation, the model traverses this process in reverse, progressively denoising latent embeddings and decoding them into RGB space. The cascaded diffusion pipeline comprises three stages. **i)** Prompt creation constructs masked, spatially aligned latents that encode the known region and the shape of the missing region. **j)** Low-resolution latent diffusion then generates a coarse embedding of the unknown region using cross-attention and U-Net–style down- and up-sampling blocks. **k)** Finally, a high-resolution RGB refinement stage denoises the coarse output to produce pixel-level embeddings that are decoded into predicted tissue cell types.

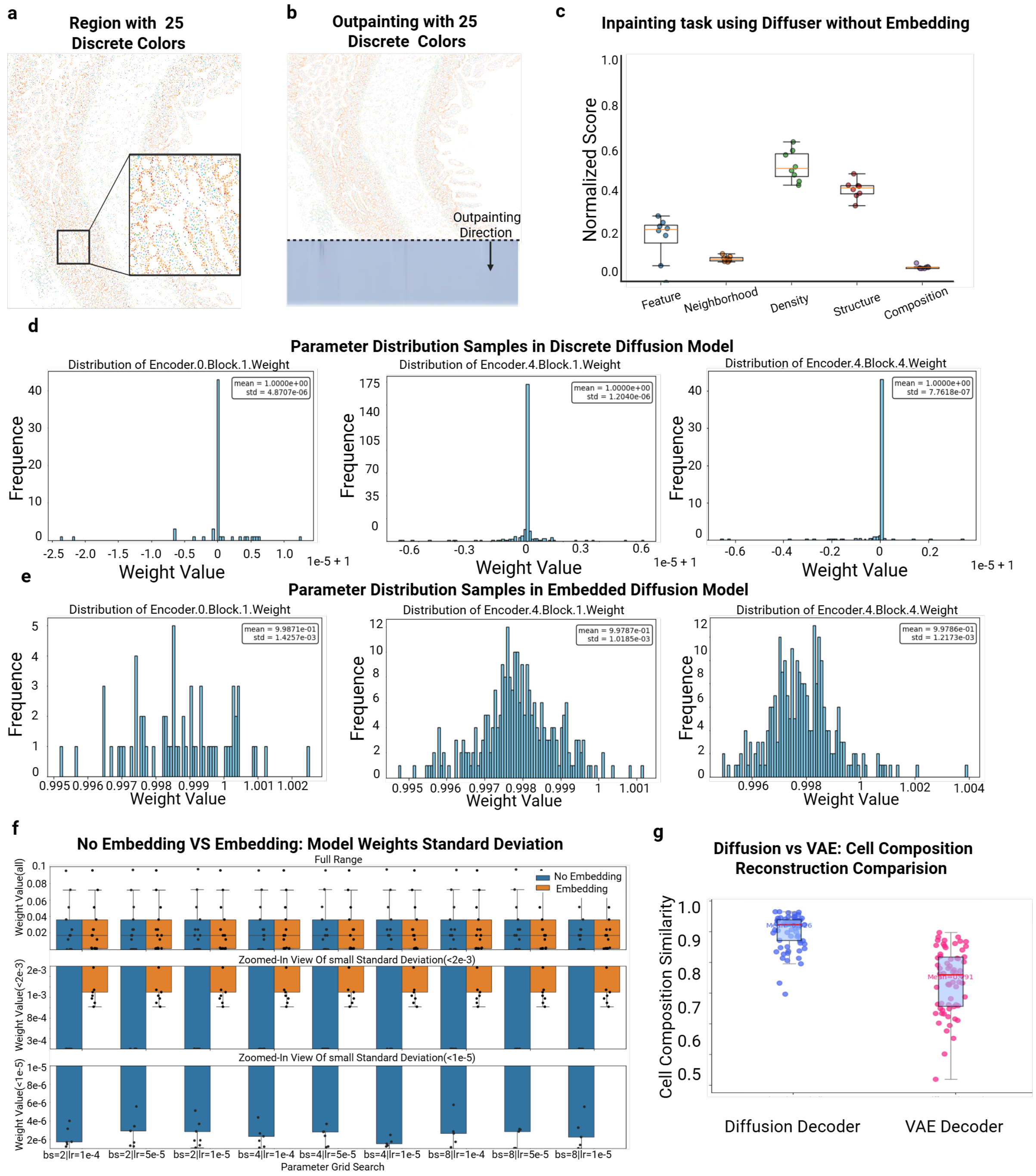

**Extended Data Fig 2** Pattern collapse happens in encoder layers without cell embedding. **a)** A sample from the reduced dataset without embedding. **b)** A generated sample from the reduced dataset without embedding where mode collapse happened. **c)** Evaluation results of SD2 inpainted tissue without embedding with 8 tissue regions, showing very low quality. **d)** Example histograms of representative encoder weights (non-embedded) showing an abnormally narrow, sharply peaked distribution. **e)** Example histograms of the same encoder layer from the embedded model showing a substantially broader, approximately Gaussian-like distribution. **f)** Layer-wise diffusion's encoder weight standard deviations for models with and without the learned embedding across 3×3 grid search: blue bars correspond to models without embedding and orange bars to models with embedding. Two zoomed-in insets highlight extremely small standard deviations (on the order of $10^{-6}$ and $10^{-5}$) observed in the non-embedded case, showing near-zero variance in many encoder layers. **g)** Comparison of reconstructing the cell compositions across CODEX intestine dataset between Diffusion decoder and VAE decoder.

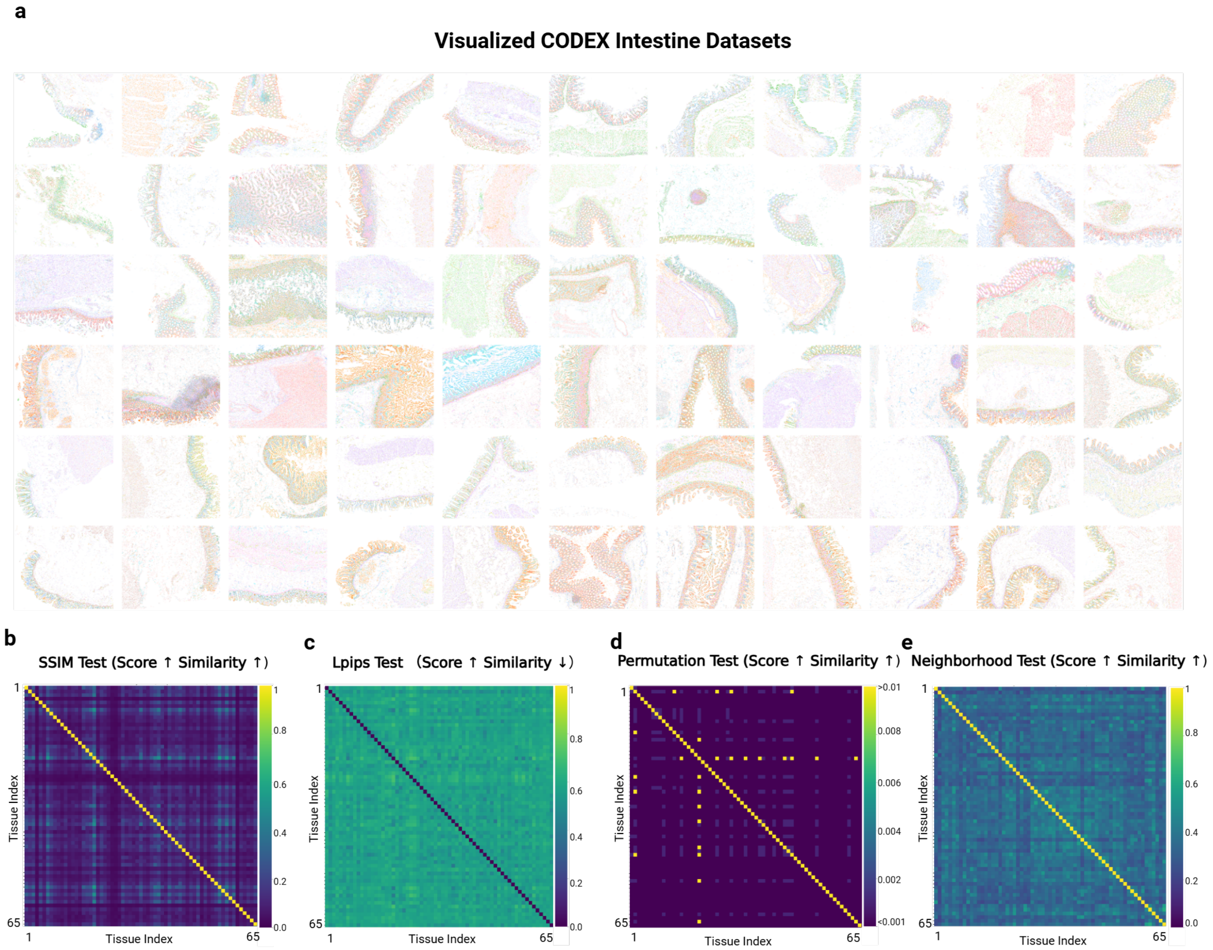


**Extended Data Fig 3** Visualization of the CODEX Intestine dataset and similarity check across dataset for validating MORPHE's generalizability. **a)** Visualization of the CODEX Intestine dataset, which comprises 66 different slices. **b)** SSIM Test (higher score indicates higher similarity) — fine-scale architectural similarity between CODEX intestine regions. **c)** LPIPS Test (higher score indicates lower similarity) — perceptual similarity of high-level texture and structure. **d)** Permutation Test (higher score indicates higher similarity) — statistical comparison of global cell-type compositions. **e)** Neighborhood Test (higher score indicates higher similarity) — local cellular-context similarity quantified by complement Jensen–Shannon divergence (JSD).

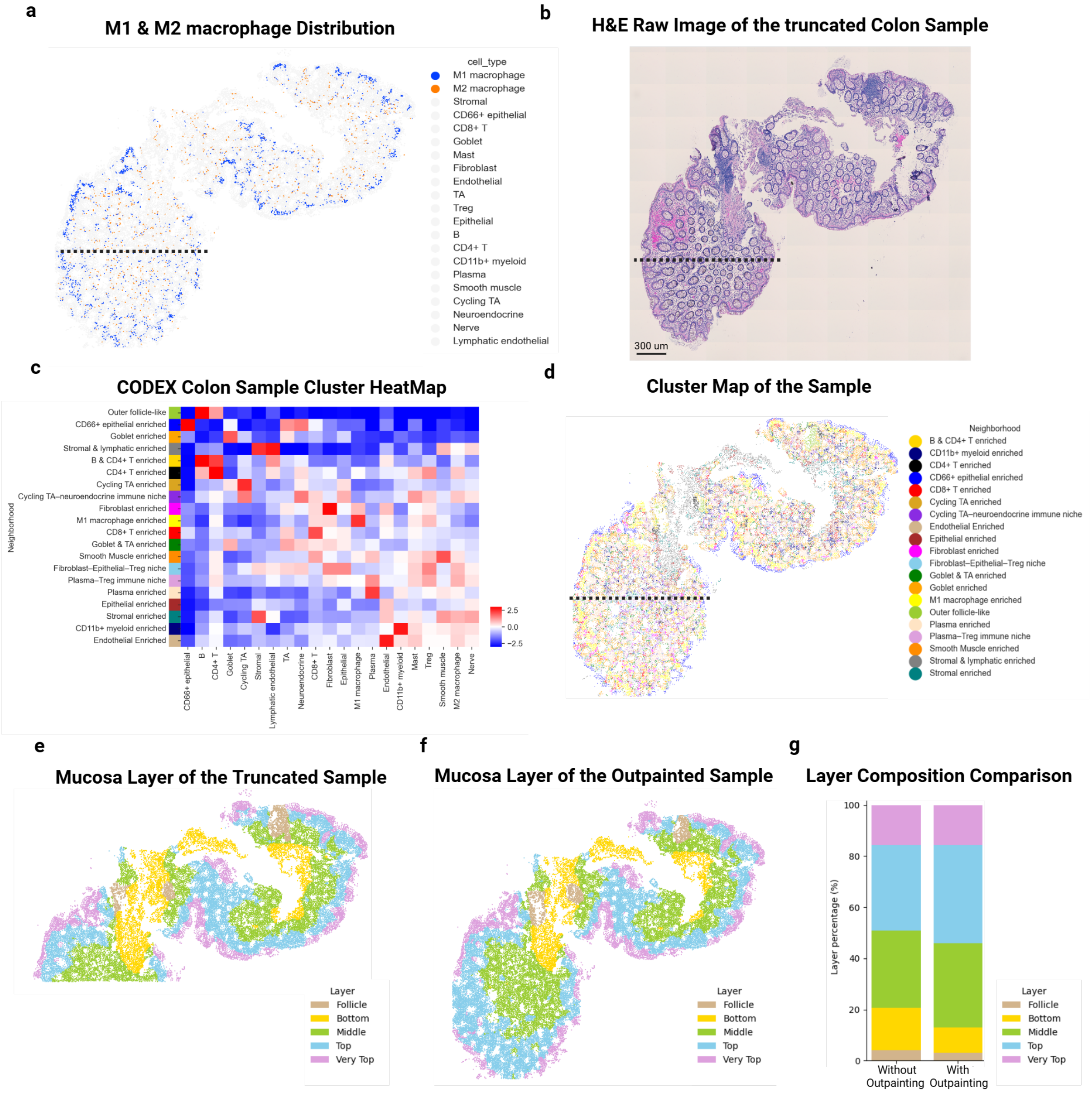


**Extended Data Fig 4 a)** M1 and M2 macrophage distribution across the whole outpainted region. **b)**Raw H&E map of the truncated sample which needs to be outpainted. **c)**Neighborhood cluter heatmap of the outpainted sample (**Fig. 3g**), which is clustering with 10 nearest neighbors. **d)**Cluter map of the outpainted sample, each cell is replaced by assigned cluster in **b**. **e)** Layer map of the sample without outpainting, where the neighborhood clusters are further grouped into five higher-order spatial categories: follicle, bottom, middle, top, and very top. **f)** Layer map of the sample with outpainting, where the neighborhood clusters are further grouped into five higher-order spatial categories: follicle, bottom, middle, top, and very top. **g)** Comparison for Layer composition of with/without outpainting sample.

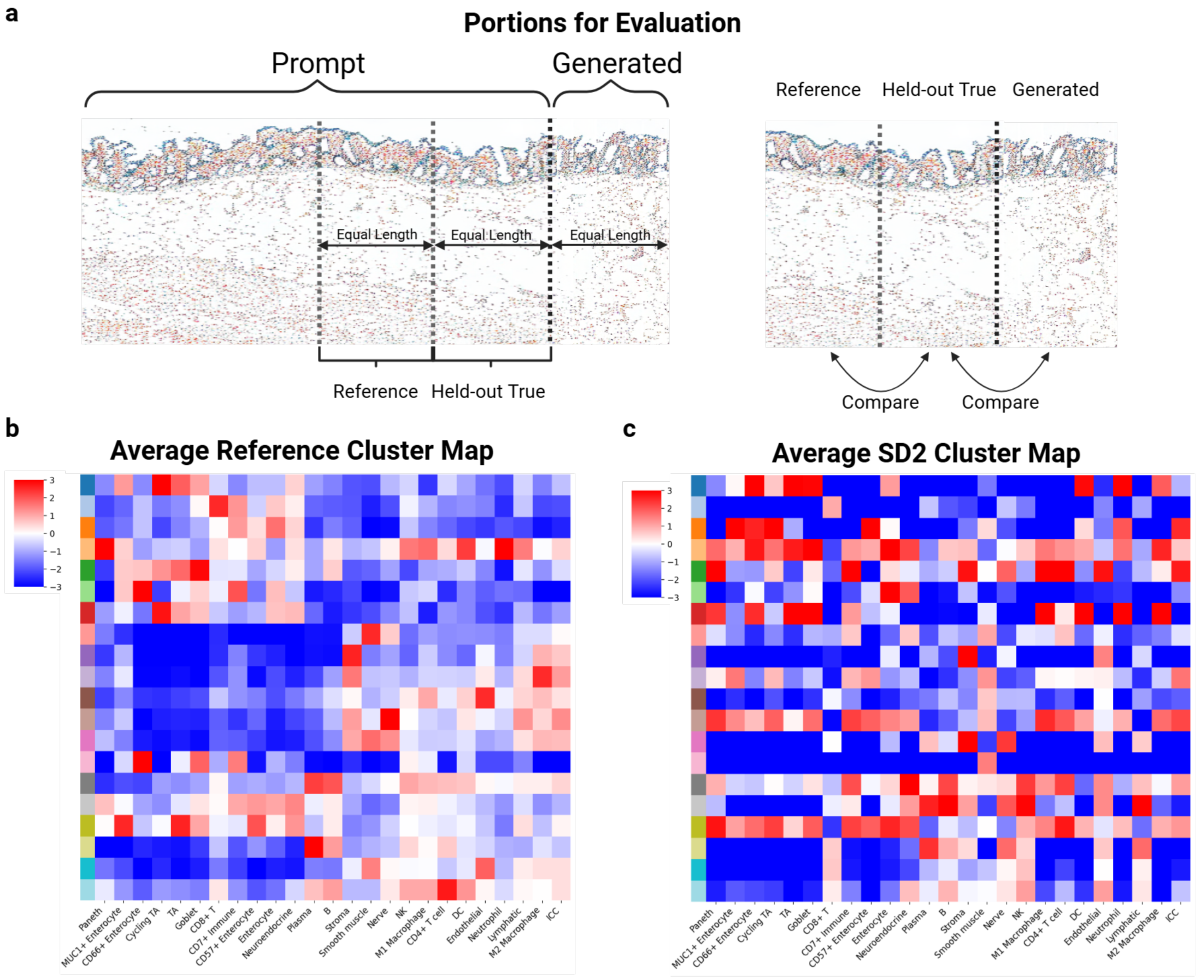


**Extended Data Fig 5 a)** Visualization of the partition we evaluated for outpainting task: reference part, held-out true part and generated part. **b)** Average neighborhood cluster heatmap shows the held-out true tissue's neighborhood in each of the clusters across the whole 66 reference regions. **c)** Average neighborhood cluster heatmap shows the SD2 generated tissue's neighborhood in each of the clusters across the whole 66 reference regions.

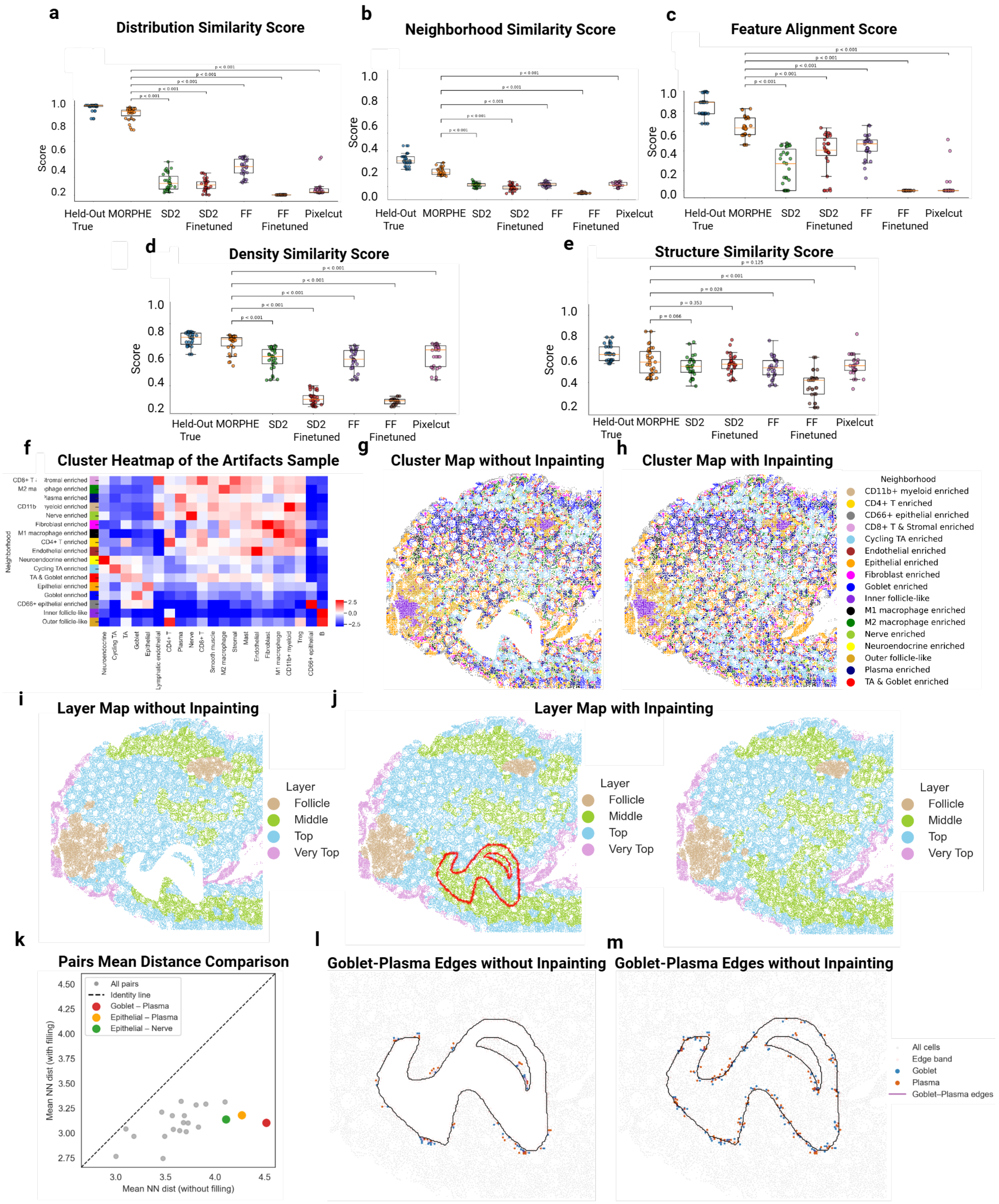


**Extended Data Fig 6** Inpainting evaluation for 5% masked area across 31 CODEX intestine regions with significance test (T-test). **a)** Distribution Similarity Score—MORPHE outperforms other state-of-the-art methods, showing significantly higher agreement between predicted and true cell-type composition distributions ($p < 0.001$). **b)** Neighborhood Similarity Score—MORPHE achieves superior local spatial consistency of reconstructed cellular neighborhoods compared with competing models ($p < 0.001$). **c)** Feature Alignment Score—MORPHE exhibits markedly better per-cell-type color alignment in RGB feature space than other models ($p <0.001$). **d)** Density Similarity Score—MORPHE more accurately recovers spatial cell-density patterns relative to baselines ($p < 0.001$). **e)**

Structure Similarity Score—MORPHE maintains comparable global tissue organization and morphology ($p = 0.066$). **f)** Neighborhood cluter heatmap of the inpainted sample (**Fig. 5g**) , which is clustering with 10 nearest neighbors. **g)** Cluter map of the sample without inpainting, each cell is replaced by assigned cluster in **f**. **h)** Cluter map of the inpainted sample, each cell is replaced by assigned cluster in **f**. **i)** Layer map of the sample without inpainting, where the neighborhood clusters are further grouped into four higher-order spatial categories: follicle, middle, top, and very top. **j)** Layer map of the inpainted sample, where the neighborhood clusters are further grouped into four higher-order spatial categories: follicle, middle, top, and very top. **k)** Pairwise mean nearest-neighbor distances with and without spatial inpainting: For each cell-type pair, the mean nearest-neighbor (NN) distance is computed using k=10 neighbors within a radius of r≤10 pix, considering only cell centers located in the edge band. The x-axis shows distances measured without inpainting, and the y-axis shows distances after inpainting. Gray points denote all cell-type pairs, while colored points highlight selected pairs (Goblet–Plasma, Epithelial–Plasma, and Epithelial–Nerve). The dashed line indicates the identity line (y=x), where inpainting has no effect. **l)** Detected Goblet–Plasma cell–cell edges within the edge band, illustrating sparse and discontinuous spatial contacts in the absence of inpainting. **m)** Detected Goblet–Plasma cell–cell edges within the edge band, illustrating more dense spatial contacts with inpainting.

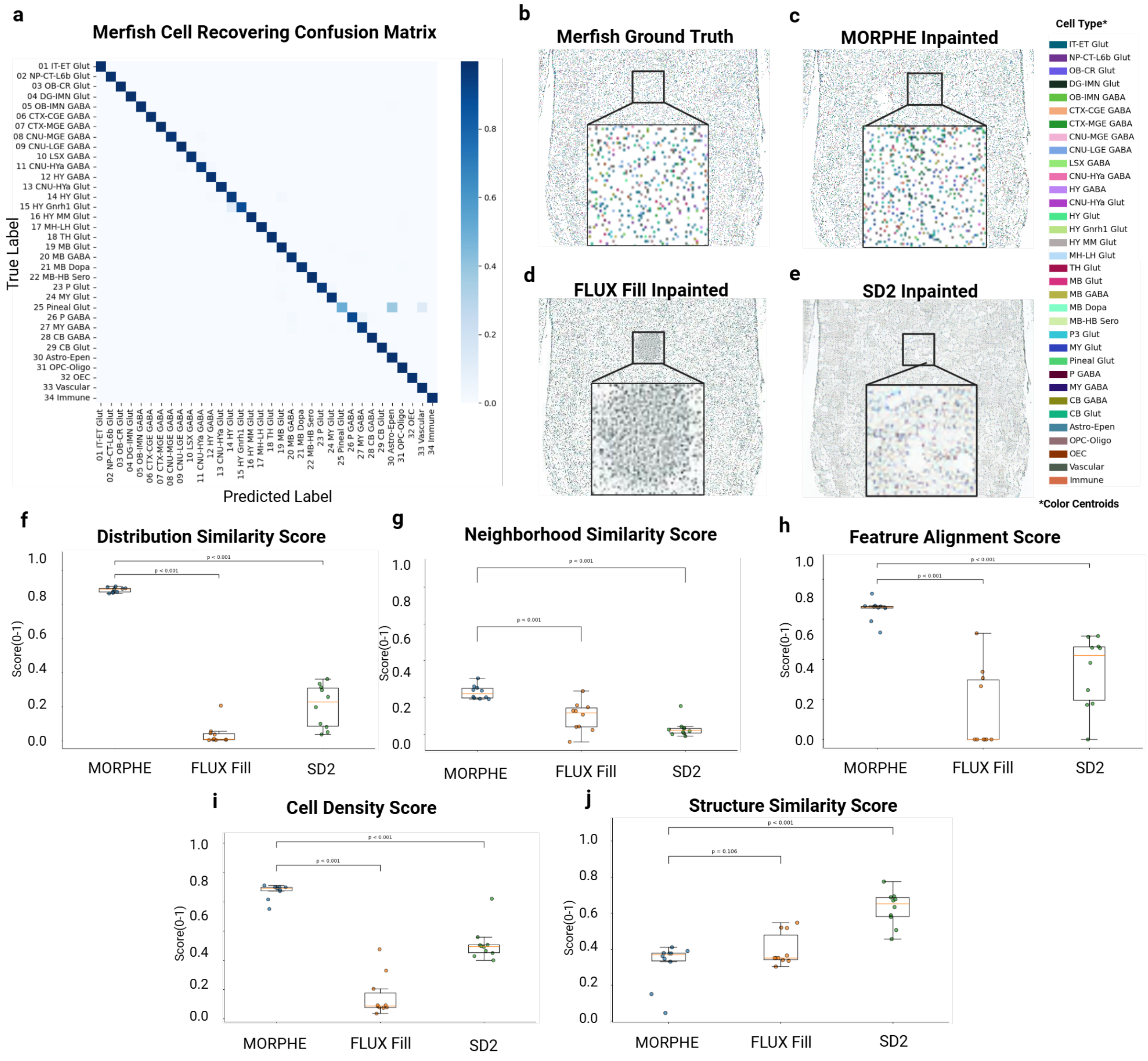


**Extended Data Fig 7** MERFISH cell recovery and inpainting evaluation. **a)** Confusion matrix showing accurate recovery of 34 MERFISH cell types by MORPHE, with strong diagonal dominance indicating high type-specific fidelity. **b–e)** Example MERFISH tissue region inpainting: (**b**) ground-truth, (**c**) MORPHE inpainted, (**d**) FLUX Fill inpainted, and (**e**) SD2 inpainted. **f–j)** Quantitative evaluation for inpainting task of 5% masked area with significance test (T-test) across 10 tissue regions: (**f**) Feature Alignment Score—MORPHE significantly outperforms other models in aligning per-cell-type features in RGB space ($p < 0.001$). (**g**) Structure Similarity Score—MORPHE achieves comparable global tissue structural consistency. (**h**) Distribution Similarity Score—MORPHE demonstrates markedly improved agreement between predicted and true cell-type composition distributions ($p < 0.001$). (**i**) Cell Density Score—our model more accurately recovers spatial cell-density patterns compared with baselines ($p < 0.001$). (**j**) Neighborhood Similarity Score—MORPHE achieves superior preservation of local microenvironmental organization ($p < 0.001$).

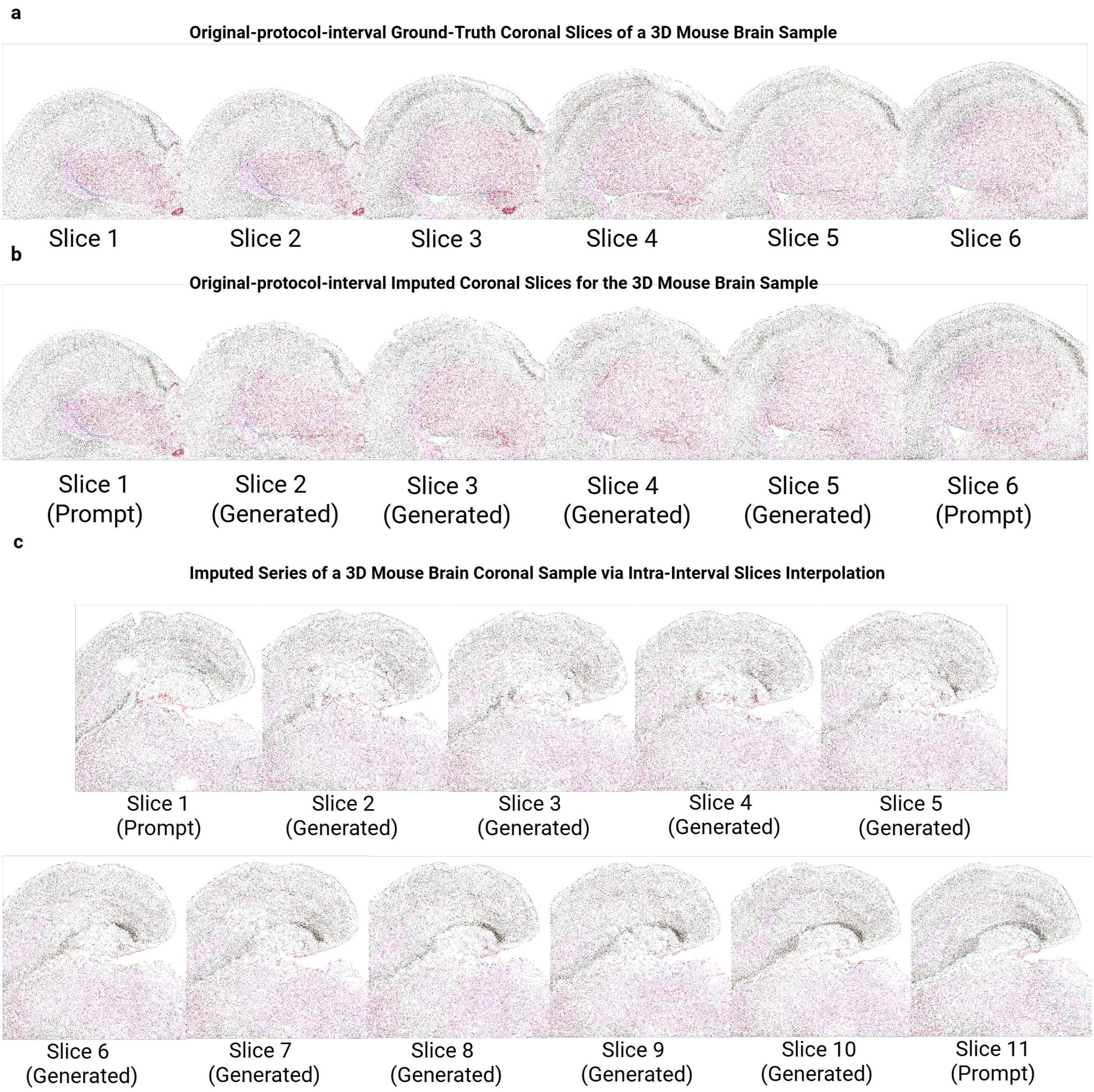


**Extended Data Fig 8 a)** larger version of Fig. 5e. **b)** larger version of Fig. 5f. **c)** larger version of Fig. 5j.

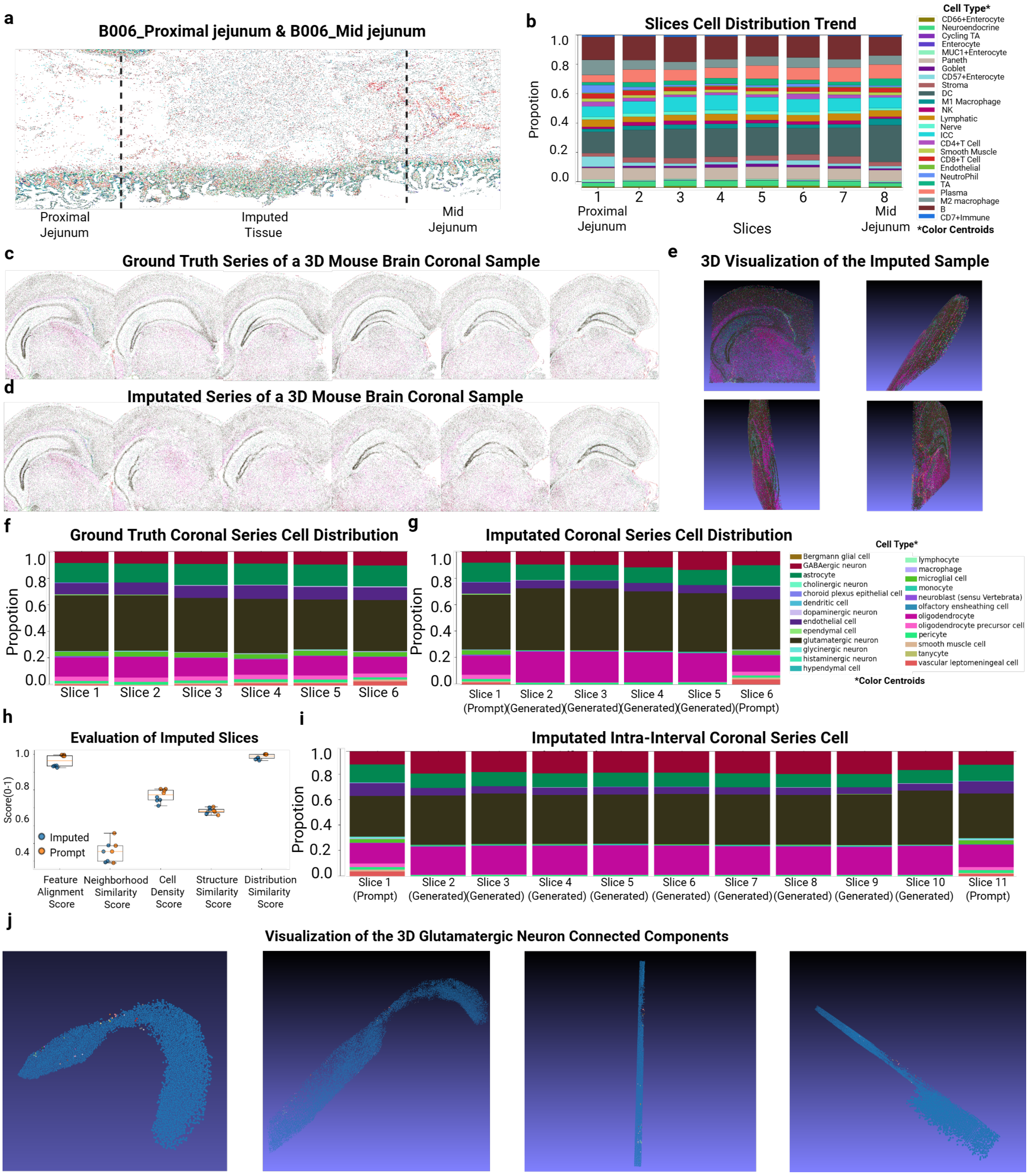


**Extended Data Fig 9** Additional examples of imputation. **a)** MORPHE fills between regions Proximal Jejunum and mid jejunum of patient B006 in CODEX dataset **b)** Cell type distribution for each slice of tissue between the two-sided prompt: B006 Proximal Jejunum and B006 mid jejunum shows the stability of cell composition. **c)** Ground truth of an original-protocol fixed-interval slices series in 3D MERFISH mouse brain data. **d)** round truth of an original-protocol fixed-interval slices series in 3D MERFISH mouse brain data. **e)** 3D visualization of the imputation slices series in **d**. **f)** Cell-type distribution of the ground truth slice series corresponding to **c**. **g)** Cell-type distribution of the imputed slice series corresponding to **d**. **h)** Metric-wise comparison between MORPHE imputed and prompt-based scores across five evaluation metrics. Blue points represent scores computed by comparing each imputed slice with its corresponding ground-truth slice. Orange points represent scores obtained by comparing the real endpoint slice (used as the prompt/input) with each intermediate ground-truth slice. This comparison reflects the intrinsic similarity between two real slices at different temporal positions, and serves as a reference baseline for interpreting the similarity scores between imputed slices and their corresponding ground-truth slices. Boxplots summarize the aggregated distribution of scores for each metric. **i)** Cell-type distribution of the imputed slice series corresponding to **Fig. 5k**. **j)** 3D visualization of the 3D connected components of the glutamatergic neuron enriched region in **Fig. 5l**, different colors indicate different 3D connected components.

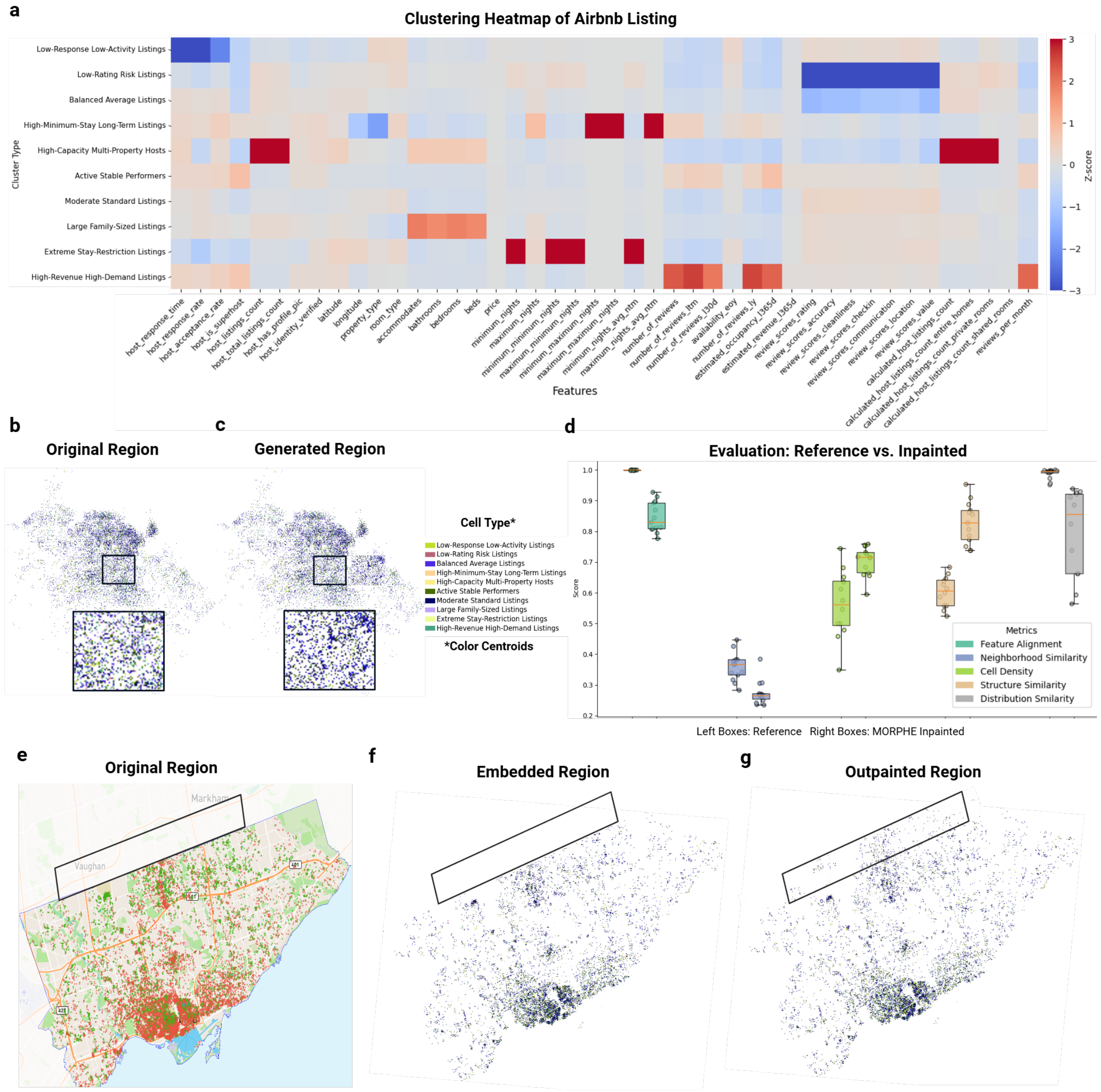


**Extended Data Fig 10** Appling MORPHE to Airbnb listings map. **a)** Clustering Airbnb listings into 10 clusters by KMeans which serves as the classification types here. Each of the type has an unique property: 0: "Low-Response Low-Activity Listings", 1: "Low-Rating Risk Listings", 2: "Balanced Average Listings", 3: "High-Minimum-Stay Long-Term Listings", 4: "High-Capacity Multi-Property Hosts", 5: "Active Stable Performers", 6: "Moderate Standard Listings", 7: "Large Family-Sized Listings", 8: "Extreme Stay-Restriction Listings", 9: "High-Revenue High-Demand Listings". **b)** One example region of Barcelona. **c)** The example region in **b)** with the mask region inpainted for quality check. **d)** Evaluation results across 16 regions using the same 5 metrics we used above to evaluate the quality on the inpainting task. To give a reference for comparison, we took a same size&shape patch near the masked area and compare it with the masked area. The left boxes of each metic are references and the right boxes are inpainted area by MORPHE. **e)** Original listing map of Toronto. **f)** Embedded listing map of Toronto. **g)** Outpainted listing map of Toronto.
